# Comprehensive multiomic profiling of somatic mutations in malformations of cortical development

**DOI:** 10.1101/2022.04.07.487401

**Authors:** Changuk Chung, Xiaoxu Yang, Taejeong Bae, Keng Ioi Vong, Swapnil Mittal, Catharina Donkels, H. Westley Phillips, Ashley P. L. Marsh, Martin W. Breuss, Laurel L. Ball, Camila Araújo Bernardino Garcia, Renee D. George, Jing Gu, Mingchu Xu, Chelsea Barrows, Kiely N. James, Valentina Stanley, Anna Nidhiry, Sami Khoury, Gabrielle Howe, Emily Riley, Xin Xu, Brett Copeland, Yifan Wang, Se Hoon Kim, Hoon-Chul Kang, Andreas Schulze-Bonhage, Carola A. Haas, Horst Urbach, Marco Prinz, Corrine Gardner, Christina A. Gurnett, Shifteh Sattar, Mark Nespeca, David D. Gonda, Katsumi Imai, Yukitoshi Takahashi, Robert Chen, Jin-Wu Tsai, Valerio Conti, Renzo Guerrini, Orrin Devinsky, Wilson A. Silva, Helio R. Machado, Gary W. Mathern, Alexej Abyzov, Sara Baldassari, Stéphanie Baulac, Focal Cortical Dysplasia Neurogenetics Consortium, Brain Somatic Mosaicism Network, Joseph G. Gleeson

## Abstract

Malformations of cortical development (MCD) are neurological conditions displaying focal disruption of cortical architecture and cellular organization arising during embryogenesis, largely from somatic mosaic mutations. Identifying the genetic causes of MCD has been a challenge, as mutations remain at low allelic fractions in brain tissue resected to treat epilepsy. Here, we report a genetic atlas from 317 brain resections, identifying 69 mutated genes through intensive profiling of somatic mutations, combining whole-exome and targeted-amplicon sequencing with functional validation and single-cell sequencing. Genotype-phenotype correlation analysis elucidated specific MCD gene sets associating distinct pathophysiological and clinical phenotypes. The unique spatiotemporal expression patterns identified by comparing single-nucleus transcriptional sequences of mutated genes in control and patient brains implicate critical roles in excitatory neurogenic pools during brain development, and in promoting neuronal hyperexcitability after birth.

## Introduction

MCDs are heterogeneous groups of neurodevelopmental disorders with localized malformation of cortical structures, often presenting with intractable epilepsy^1^. Major MCD subtypes include different classes of focal cortical dysplasia (FCD), hemimegalencephaly (HME), and tuberous sclerosis complex (TSC)^2^. The International League Against Epilepsy (ILAE) has classified FCD subtypes based on neuropathological features and cell types^3^. MCD patients often undergo surgical resection of the lesion to treat drug-refractory epilepsy, which has led to remarkable clinical benefits in published series^4^. The abnormal histology of resected regions includes loss of lamination of cortical layers, enlarged dysplastic neurons, or balloon cells, sometimes accompanied by other brain abnormalities. But similar to brain tumors, it can be difficult to predict pathology prior to surgery.

Again, like with brain tumors, genetic studies may offer insights into mechanisms. Somatic mTOR pathway gene mutations are frequently detected in HME and type II FCD foci^5,6^. Recently, small- or medium-size cohort studies (<100 cases) have confirmed these results and have correlated defects in neuronal migration, cell size, and neurophysiology^7^. Still, the vast majority of MCD cases still remain genetically unsolved, suggesting other genes or modules contribute to MCD.

Detecting mutant alleles in bulk resected foci from MCD patients is challenging because unlike in brain tumors, the mutant cells in MCD are probably not hyperproliferative, and thus variant allelic fraction (VAF) are often <5%, diluted by genomes of surrounding non-mutated cells^8^. Fortunately, new computational algorithms have helped reduce false-positive and false-negative signals, even when no ‘normal’ paired sample is available for comparison^9-11^. The NIH-supported Brain Somatic Mosaicism Network established the ‘BSMN common pipeline’, incorporating a ‘best practice’ workflow to reliably and reproducibly identify somatic variants contributed by members of the Network^12^. With these advances, we thus assessed the possibility of gene networks beyond mTOR that could underlie MCDs. This new gene discovery may give insights into novel druggable pathways in cases of incomplete resection due to regional importance or drug-resistant forms of MCD.

## Results

### The genetic landscape of MCD from targeted and unbiased sequencing

To perform a thorough genetic screening of somatic mutations in resected epileptic tissue, we formed the FCD Neurogenetics Consortium and enrolled 327 samples that met clinical and pathological criteria for FCD or HME. We excluded TSC from our enrollment criteria because genes are already well known. Our cohort included 31 HME cases, 98 type I-, 142 type II-, 32 type III-, and 12 unclassified-FCD cases. We included acute resected brains from 10 neurotypicals and 2 TSC cases for comparison (Fig. 1a, supplementary table 1). Patients with environmental causes, syndromic presentations, inherited mutations, multifocal lesions, or tumors were excluded (Methods).

**Figure 1.**
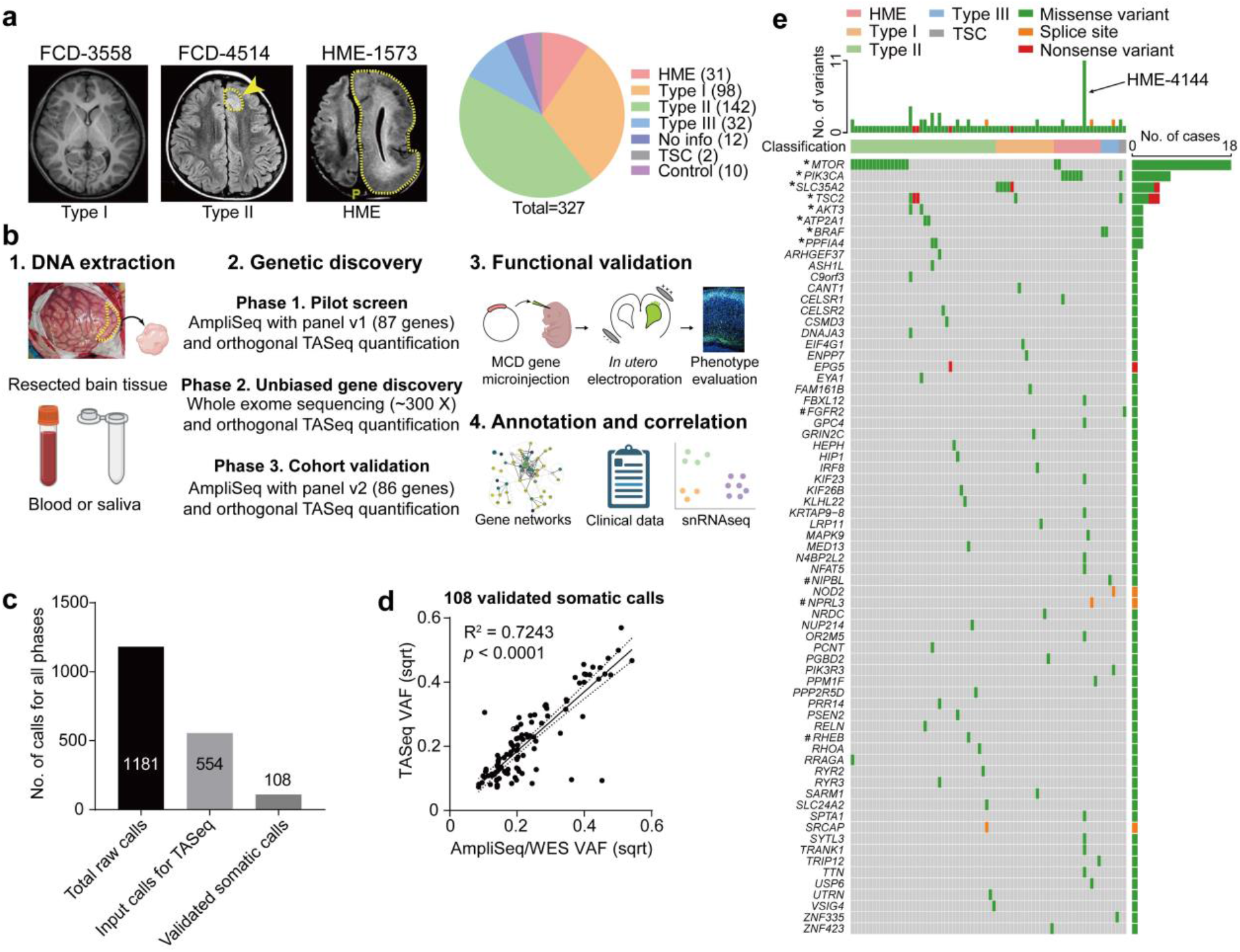
Comprehensive genetic profiling and validation of somatic variants in 327 MCD patients. (a) Representative MRI image of FCD-3588 (FCD type I), FCD-4514 (FCD type II), HME-1573, and a pie chart for the composition of our MCD cohort. Yellow arrow and dash: affected brain regions. (b) Workflow for comprehensive genetic profiling of MCD, using a three-phase approach from patient DNA. Each phase was followed by quantification/validation of each variant with target amplicon sequencing (TASeq). Phase 1] 1000 × pilot screening of DNA with an 87-gene mTOR-related panel. Phase 2] 300 × whole-exome sequencing (WES) with best-practice somatic variant discovery for novel candidate discovery. Phase 3] Cohort-level validation with an updated, high-confidence TASeq gene set based on knowledge from Phase 1 and 2. A subset of the somatic mutations was further functionally validated by mouse modeling. Candidate genes were annotated and correlated with external datasets such as STRING DB, clinical phenotype dataset, and newly generated single-nucleus RNAseq dataset from MCD brain. (c) Somatic variant calls were detected from all three phases of genetic discovery, yielding 108 validated somatic calls. (d) Correlation between square-root transformed (sqrt) AmpliSeq/WES variant allele fraction (VAF) and TASeq VAF. Solid line: best-fit line linear regression. Dotted lines: 95 % confidence band of the best-fit line. (e) Oncoplot with all 69 validated somatic SNVs from this study. Top: most patients had one gene mutated, a few patients had more than one gene mutated, and patient HME-4144 had 11 different validated gene mutations. Color: type of variant. * and #: recurrent genes in our cohort, and non-recurrent in our cohort but recurrent in other studies, respectively.

We used a three-phase genetic screening, each followed by filtering for likely causative mutations using published methods^13,14^, and each followed by orthogonal targeted amplicon sequencing (TASeq) intra-case validation and VAF quantification compared with controls (∼5000 X, TASeq)(Fig. 1b). In Phase 1, we performed amplicon sequencing (AmpliSeq, ∼1000 X) profiling the entire open reading frame of 87 genes previously detected in FCD/HMEs or known PI3K-AKT3-mTOR interactors (‘MCD panel v1’, Supplementary Table 2a). In Phase 2, for 75 unsolved cases from Phase 1 and additionally collected 54 cases, we performed unbiased deep whole-exome sequencing (WES, ∼300 X) on paired samples, where available, or on unpaired samples (i.e. brain plus blood/saliva vs. brain only). In Phase 3, from an additional subcohort of 132 new cases, we designed the ‘MCD panel v2’ (Supplementary Table 2b) including known and novel genes detected in Phases 1 and 2 (Extended Data Fig. 1, Methods). We re-sequenced unsolved cases from Phase 2, expecting that the higher read depth afforded by panel sequencing could provide greater sensitivity to detect low VAF mutations, and used BSMN best practice guidelines for mapping and variant calling^12^.

From Phases 1 to 3, 1181 candidate somatic SNVs were identified. Of these, 628 were excluded based on gnomAD allele frequencies, dinucleotide repeats, homopolymers, and additional BSMN established criteria (Methods)^15,16^. This yielded 554 candidate somatic SNV that were further assessed by TASeq, yielding 108 validated somatic SNV calls (19.4% validation rate, Fig. 1c, Supplementary Table 3), compared to other BSMN effort validation rates in WGS^12,17^. In detail, 15, 67, and 26 validated somatic SNV calls were derivated from phase1, 2, and phase 3, respectively. The measured VAFs between the AmpliSeq/WES and TASeq were correlated as expected (R^2^= 0.7243) (Fig. 1d). Of the 69 candidate MCD genes mutated in 76 patients, 8 were recurrently mutated, including known mTOR pathway genes as well as several novel candidates (Fig. 1e).

We estimate only ∼7% of mutations identified are likely attributable to false discovery during variant calling, based upon background mutation rate in 75 BSMN neurotypical brain samples, and published experience from the BSMN^12,18^, processed with the same workflow (see Methods). Thus, 93% of our candidate MCD mutations would not have been identified in a size-matched neurotypical control cohort.

Most patients (80.52%, 62 cases) showed a single somatic mutation, but some showed two somatic mutations (14.29%, 11 cases), and some showed more than two mutations (5.19%, 4 cases). Interestingly, HME-4144 showed 11 different somatic mutations, all of which were validated with TASeq. Although there are several possible explanations for HME-4144, we expect this reflects clonal expansion from a driver mutation, with detection of multiple passenger mutations, as reported in brain tumors^19^.

Single-base mutational signatures (SBS) were developed to describe potential mutational mechanisms in human disease^20^. We found 60.2% of mutations were C>T, likely arising from DNA epigenetic marks^21^ (Extended Data Fig. 2). Enrichment of SBS1 and SBS5, clock-like mutational signatures suggest endogenous mutations arising during corticogenesis DNA replication.

### Functional dissection of the MCD genes

Interestingly, most validated genes were non-recurrently mutated (88.4%, 61 of 69) in our cohort, suggesting substantial genetic heterogeneity in MCD. This nevertheless provided an opportunity to study converging functional gene networks. Thus, we performed Markov clustering with a STRING network generated from the putative MCD genes^22^, as well as recently reported novel MCD candidates (*NAV2, EEF2, CASK, NF1, KRAS, PTPN11*)^23,24^ (Fig. 2a). We identified four clusters, with cluster 1 (“mTOR pathway”) showing the highest term enrichment to the mTOR/MAP kinase signaling, supporting prior results for Type II MCDs. Cluster 1 also highlighted newly identified genes *FGFR2, KLHL22, RRAGA, PPP2R5D, PIK3R3, EEF2, EIF4G1*, and *MAPK9*. Cluster 2 identified “Calcium Dynamics” and included genes *ATP2A1, RYR2, RYR3, PSEN2, TTN, UTRN*. Cluster 3 was labeled “Synaptic Functions” and included genes *CASK, GRIN2C, and PPFIA4*. Cluster 4 was labeled “Gene Expression” and included intellectual disability genes, mostly involved in nuclear function, including *NUP214, PRR14, PCNT, NIPBL, SRCAP, ASH1L, TRIP12*, and *MED13* (Fig. 2b).

**Figure 2.**
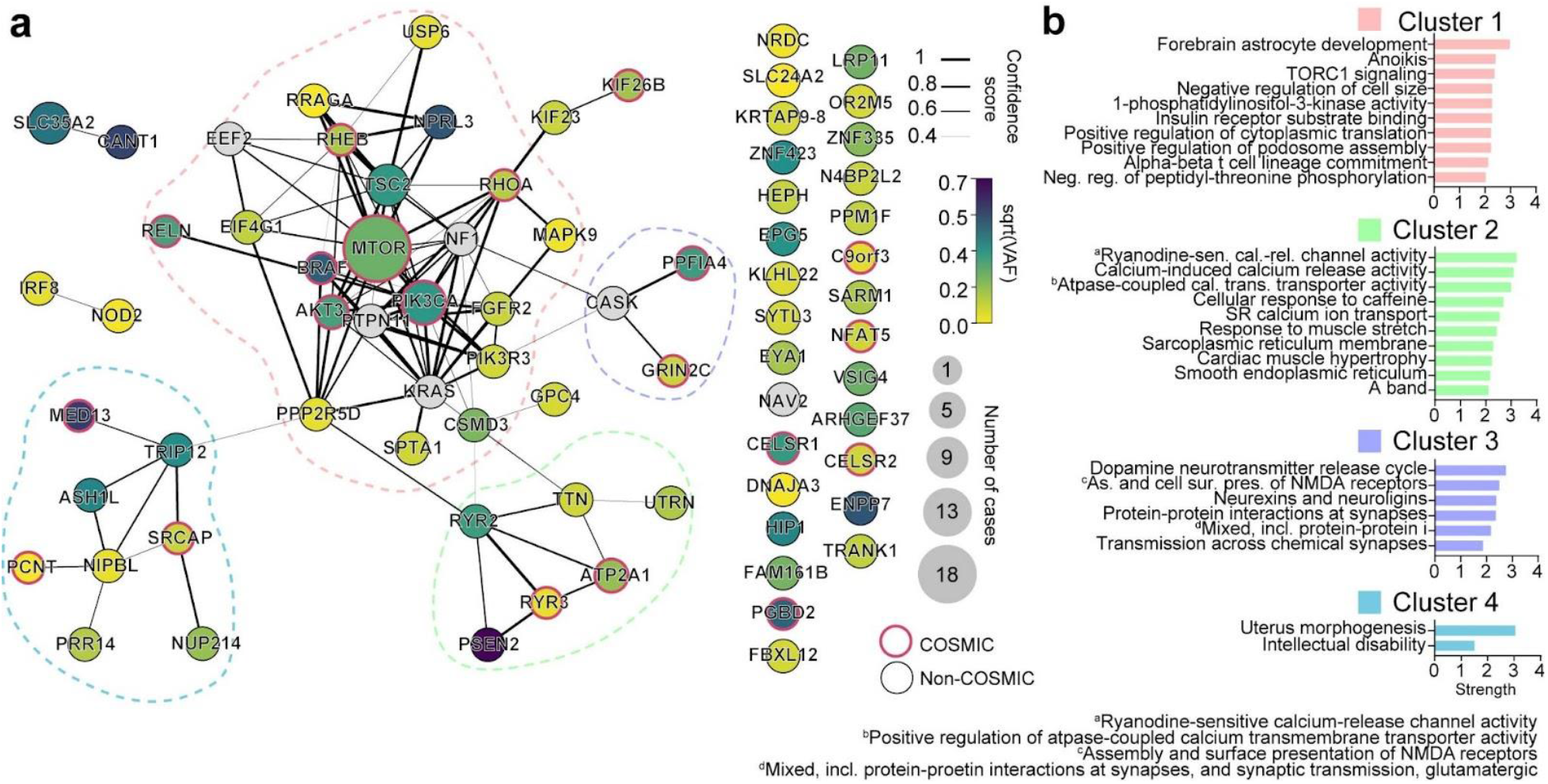
Four major gene networks were discovered from the comprehensive MCD gene profiling. (a) STRING DB pathway analysis of the 69 MCD discovered genes and six novel genes from recent publications identifies MTOR/MAP kinase pathway (pink, Cluster 1), Calcium dynamics (green, Cluster 2), Synapse (purple, Cluster 3), Gene expression (blue, Cluster 4). Edge thickness: confidence score calculated by STRING. Size and color of a node: square root transformed (sqrt) number of patients carrying a given mutation and average sqrt VAF across all patients, respectively. Non-clustered orphan genes are listed on the right. Red border: variant reported in the COSMIC database. (b) Gene Ontology (GO) analysis results confirmed the functions of compositions in each network. Top GO terms or KEGG pathways. Strength calculated by STRING.

Notably, *ATP2A1, PPFIA4*, and *NIPBL* were recurrently mutated, either within our cohort or with a recent report^24^ (Extended Data Fig. 3a-b), occurring within the latter 3 clusters. While these clusters were not previously reported in MCDs, they were previously implicated in epilepsy, neurodevelopmental and neurodegenerative disease^25,26^, suggesting functional overlap with MCDs. We further performed ClueGO analysis and found enrichment in mTOR signaling, focal adhesion assembly, cardiac muscle cell contraction, and artery morphogenesis (Extended Data Fig. 4). ClueGO also displayed isolated gene ontology (GO) term clusters such as ‘calcium ion import’ and ‘protein localization to synapse’.

### Functional validation of selected module genes in embryonic mouse brain

To investigate the roles of novel MCD genes and modules, we selected two potential mTOR pathway mutations (*RRAGA* p.H226R, *KLHL22* p.R38Q), and non-mTOR gene mutation (*GRIN2C* p.T529M), discovered in FCD-7967, 3560, and 5157, respectively. *RRAGA* encodes Ras-related GTP binding A (RAGA), a GTPase sensing amino acid and activating mTOR signaling, with two functional domains: GTPase domain and C-terminal ‘roadblock’ domain (CRD) ^27^. The mosaic p.H226R mutation occurs within the CRD, which binds to the RAGB protein and is conserved throughout vertebrate evolution (Extended Data Fig. 3c) and thus could change binding affinity. *KLHL22* encodes a CUL3 adaptor, determining E3 ubiquitin ligase specificity. The CUL3-KLHL22 complex mediates the degradation of DEPDC5, required for mTORC1 activation^28^. The KLHL22 *p*.R38Q variant in FCD-3560 is near the BTB (Broad-Complex, Tramtrack, and Bric-à-brac) domain that interacts with CUL3 (Extended Data Fig. 3d), suggesting the variant could enhance mTORC1 activity. *GRIN2C* encodes a subunit of the NMDA receptor regulating synaptic plasticity, memory, and cognition^29,30^, dysfunction of which is implicated in many neurocognitive diseases including epilepsy, neurodevelopment, and tumors^31,32^. *GRIN2C* p.T529M mutation is located in the S1 glutamate ligand-binding domain (S1 LBD) (Extended Data Fig. 3e). *GRIN2A* p.T531M mutation, an analog mutation of *GRIN2C* p.T529M in our cohort, was previously reported in epilepsy-aphasia spectrum disorders, where it increased NMDA receptors ‘open-state’ probability^32^. This suggests that the p.T529M mutation activates the channel, likely in an mTOR independent fashion. Thus, all mutations assessed here are likely gain-of-function and exert functional impact on cells in which they are expressed.

To test this hypothesis, we introduced mutant or wildtype (WT) genes co-expressing enhanced green fluorescent protein (EGFP) into the dorsal subventricular zone via electroporation at mouse embryonic day 14 (E14), then harvested tissue at either E18 to assess migration, or at postnatal day 21 (P21) to assess cell size and phospho-S6 as a reporter of mTOR activity^33^ (Fig. 3a). In E18 cortices, we found EGFP-positive cells expressing mutant but not WT forms of *RRAGA* and *KLHL22* showed significant migration defects of varying severity, whereas mutant *GRIN2C* showed no defect (Fig. 3b). These migration defects in *RRAGA* and *KLHL22* mutant cells replicate major findings of MCD disrupted cortical architecture.

**Figure 3.**
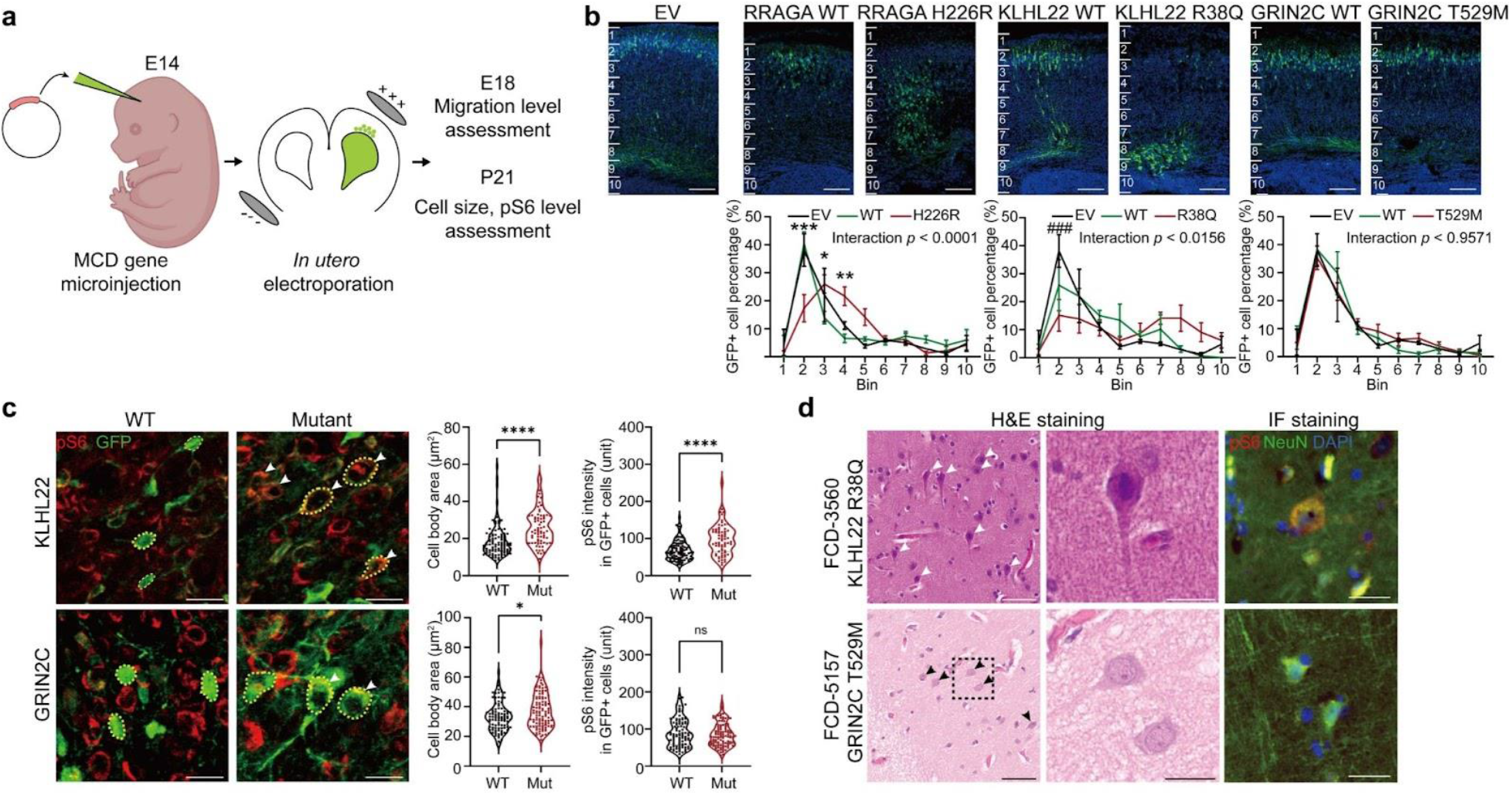
Selected novel MCD somatic variants show functional defects in embryonic mouse brain and patient samples. (a) Workflow for functional validation of candidate mosaic variants in mice. (b) Two different mutations in novel FCD type II genes, *RRAGA* H226R and *KLHL22* R38Q, but not a novel FCD type I gene, *GRIN2C*, disrupt cellular radial migration from the subventricular zone (SVZ). Below: two-way ANOVA and Sidak multiple comparisons with *p*-values of interaction between genotype and bin factor. * or # indicates a *p*-value in comparison between WT and mutant group, or EV and mutant group respectively. Ten bins from the surface of the cortex (top) to SVZ (bottom). Scale bar: 100 μm. Error bar: ±SE. (c) Immunofluorescence in postnatal day 21 mouse cortices for KLHL22 and GRIN2C wild-type (WT) or mutant isoform. Neurons expressing mutant KLHL22 and GRIN2C recapitulate histological phenotypes shown in (d), with enlarged cell bodies (white arrow) compared to WT isoforms (WT control), whereas only neurons expressing KLHL22 but not GRIN2C mutant isoform display increased pS6 levels compared to control. Dotted lines: examples of cell body size quantification. Two-sided Student’s *t*-test. Scale bar: 20 μm. (d) H&E and phospho-S6 (pS6) staining of the resected brain from FCD-3560 and FCD-5157. Box area is zoomed in the middle image. Arrows: dysplastic cells. Right: Immunofluorescence (IF) for pS6 and NeuN. Note dysplastic pS6-positive neurons with increased pS6 levels are present in FCD-3560 but not in FCD-5157. Scale bar: 60 μm on the left, 20 μm on the middle and right. ****p < 0.0001; *p < 0.05; ns, non-significant. ###p < 0.001. EV: empty vector.

We next assessed cellular phenotype at P21 with samples available in both mice and the corresponding patients and found enlarged cell body area in both mutant forms of *KLHL22* and *GRIN2C* compared to according wildtype. In contrast, the elevated levels of pS6 staining, described previously in association with mTOR pathway mutations^6^, was found only in mutant *KLHL22*, but not in mutant *GRIN2C* mice (Fig. 3c).

To assess correlation with human samples, we assessed archived neuropathological tissue sections for histology and pS6 activity. Similar to our mouse models, we found patient FCD-3560 carrying *KLHL22* p.R38Q showed enlarged neurons that co-stained for excess pS6 staining, whereas FCD-5157 carrying *GRIN2C* p.T529M showed only a slight increase in cell body size and no evidence of excessive pS6 staining (Fig. 3d). While this analysis does not take into account the genotype of individual cells, it suggests *KLHL22* but not *GRIN2C* mutations impact mTOR signaling.

### Genotype-phenotype correlations in MCD patients

To assess the phenotypic contributions of the MCD genes we found, we focused on 76 of our ‘genetically solved’ MCD cases, comparing detailed neuropathology, brain imaging, and clinical course. We performed Pearson correlation followed by hierarchical clustering based upon ILAE neuropathological diagnosis, compared with GO term-based curated genesets and whether the genetic variant was present in COSMIC DB (Methods, Supplementary Table 3,4, Fig. 4). We found that FCD Type IIA and Type IIB, and HME were more tightly clustered than FCD Type I or III (Fig. 4a), likely reflecting shared neuropathological features that include large dysplastic neurons. As expected, FCD Type IIA, Type IIB, and HME were positively associated with the mTOR pathway GO term and COSMIC DB entry, FCD Type III, however, was associated with the MAPK pathway, consistent with recent publications implicating *BRAF, FGFR2, NOD2*, and *MAPK9* in their etiology^34-36^. FCD Type I showed few strong positive correlations for glycosylation, consistent with recent findings of somatic mutations in *SLC35A2* and *CANT1*^37,38^.

**Figure 4.**
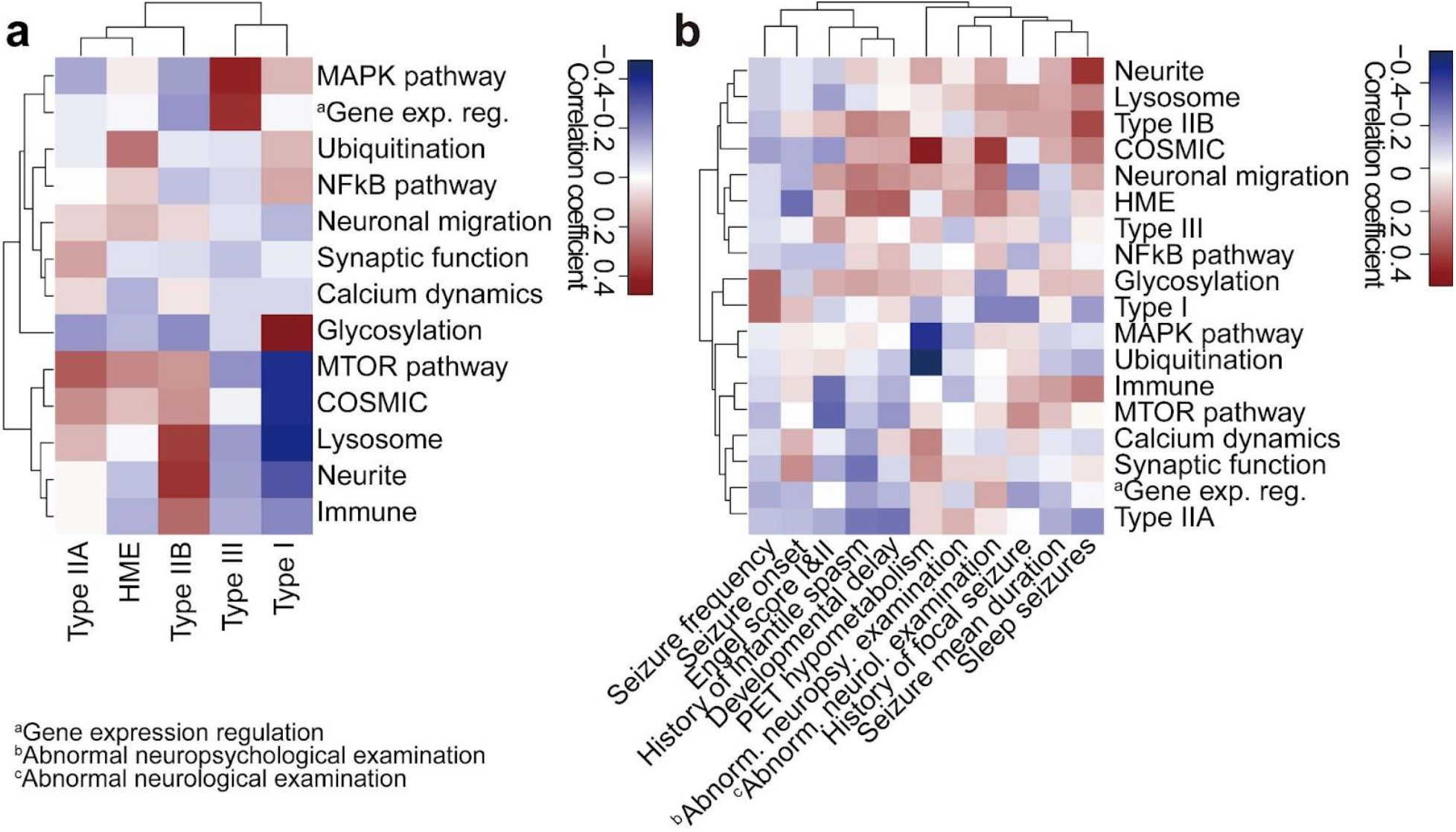
Clinical phenotypic outcomes correlate with genotype-based classifications in MCD. (a) Correlation heatmap for classification based on genetic information (y-axis) vs. International League Against Epilepsy (ILAE) classification based on histology (x-axis) using Pearson correlation. Shade: the value of Phi coefficient. Note Type IIA and HME are enriched with mTOR and Ubiquitination genes, while Type I is enriched in Glycosylation and depleted in MTOR and COSMIC genes. HME: hemimegalencephaly. (b) Correlation between classification based on genetic information and various clinical phenotypes. Shade: the value of Phi (binary data) or Pearson (continuous) correlation coefficient. For example, positron emission tomography (PET) hypometabolism is enriched in COSMIC genes and depleted in the MAPK pathway, whereas abnormal neurological examination is enriched in COSMIC genes. The whole dataset is in Supplementary Table 4.

We next investigated correlations between clinical phenotypes extracted from detailed medical records including seizure type, neuropsychological examination, and positron emission tomography (PET) metabolism, often used to help localize seizure focus^39,40^. Seizure frequency, early age of onset, Engel score, and history of infantile spasms drove clinical clustering, likely reflecting shared clinical features in the most challenging patients. Focusing on the correlations, PET hypometabolism correlated positively with COSMIC DB entry, and negatively with MAPK and Ubiquitination (Fig. 4b), suggesting divergent metabolic mechanisms. Abnormal neurological examination correlated positively with COSMIC DB entry and negatively with Type I histology, which may reflect the effects of mutations on baseline neurological function.

### MCD genes enriched in the excitatory neuronal lineage

To infer the cell type in which MCD genes function, we accessed a published single-cell transcriptome dataset from the 2nd-trimester human telencephalon, at a time when these mutations probably arose^41^ (Fig. 5a). We generated an eigengene, by mapping the average expression of our MCD genes against the UMAP plot (Fig. 5b). This showed a strong positive correlation with dividing radial glial cells, and a moderate correlation in dividing intermediate progenitor cells (IPCs) and mature excitatory neuron cells. We found a negative correlation with inhibitory neuronal lineages including medial and central ganglionic eminences (MGE, CGE) and mature interneuron clusters (Fig. 5c). We next performed deconvolution into four major module eigengene (MEs), which revealed cell types classified as mature excitatory neurons (turquoise and blue), microglia (brown), and unassigned (grey) (Fig. 5d). Quantification supported enrichment in dividing radial glia, excitatory neurons, and microglia, the latter likely driven by MCD candidate genes *IRF8* and *VSIG4* (Fig. 5e). Taken together, the expression of MCD genes is more enriched in dorsal cortex neurogenic pools and implicated in the maturation of excitatory rather than inhibitory neurogenic pools, as well as microglia.

**Figure 5.**
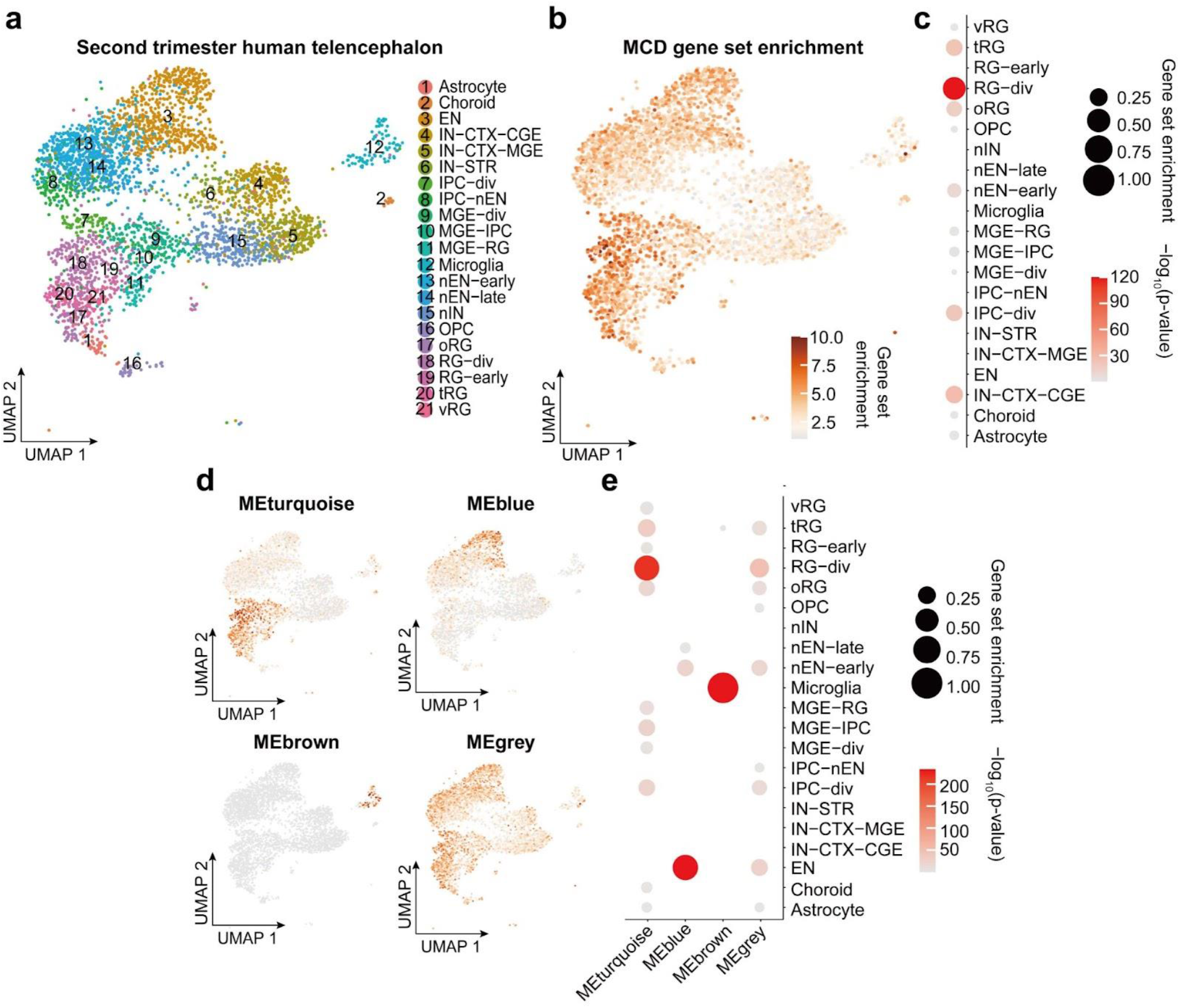
Single-nucleus transcriptomes reveal MCD gene enrichment in radial glia and excitatory neurons in the developing human cortex. (a) Uniform Manifold Approximation and Projection (UMAP) for single-nucleus transcriptome in 2nd-trimester fetal human telencephalon from a public dataset^41^. (b) UMAP enrichment patterns of an eigengene using MCD genes. Note enrichment for excitatory neurons and radial glia (dark brown). vRG: vertical radial glia, tRG: truncated radial glia, RG-div: dividing radial glia, oRG: outer radial glia, EN: excitatory neuron, nEN: newborn excitatory neuron, IPC: intermediate progenitor cell, STR: striatum, IN: interneuron, CTX: cortex, MGE: medial ganglionic eminence, CGE: central ganglionic eminence. (c) Quantification of enrichment of (b) based on cell types, showing enrichment for RG-div. (d) Four eigengenes decomposed from (b). (e) Quantification of enrichment of (d) based on cell types showing enrichment in dividing radial glia, microglia, and inhibitory cortical neurons from the medial ganglionic eminence (MGE).

### MCD gene expression is enriched in dysplastic cells

We next performed differentially expressed gene (DEG) analysis in the MCD brain. We reasoned that single-nucleus transcriptomes would be more revealing than bulk transcriptomes, but the average VAF of ∼6% in our MCD cohort meant that the vast majority of sequenced cells would be genetically wild-type. We thus decided to focus snRNAseq on resected cortex from patients with shared pathological MCD hallmarks but higher VAFs. We selected four resected brain samples, two from patients with HME (HME-4688 *PIK3CA* p.E545K, 25.1% VAF and HME-6593 *PIK3CA* p.H1047R, 13.1% VAF), and two from patients with TSC meet full diagnostic criteria. We also included brains from four neurotypical cases as a comparison and sequenced a total of 22,067 nuclei (see Methods).

While the TSC brain single nucleus transcriptomes showed substantial overlapping pools with controls, HME brains showed a distinct UMAP distribution, located at the edges of the plot (Fig. 6a). We found that very few HME cells matched expression patterns for typical brain cells, even after standard normalization and scaling (Fig. 6b, Extended Data Fig. 5a, see Methods). We thus labeled these clusters according to their closest relatives based upon established marker gene expression in the control brain, labeled as ‘astrocyte-like (Ast-L)’ or ‘oligodendrocyte-like (OD-L)’. Even with these categories, some clusters remained undefined (U) (Extended Data Fig. 5b,c). Interestingly, there was no single cell cluster that matched the VAF in the brain, suggesting the mutant cells, as well as surrounding non-mutant cells, have dramatically disrupted transcriptomes.

**Figure 6.**
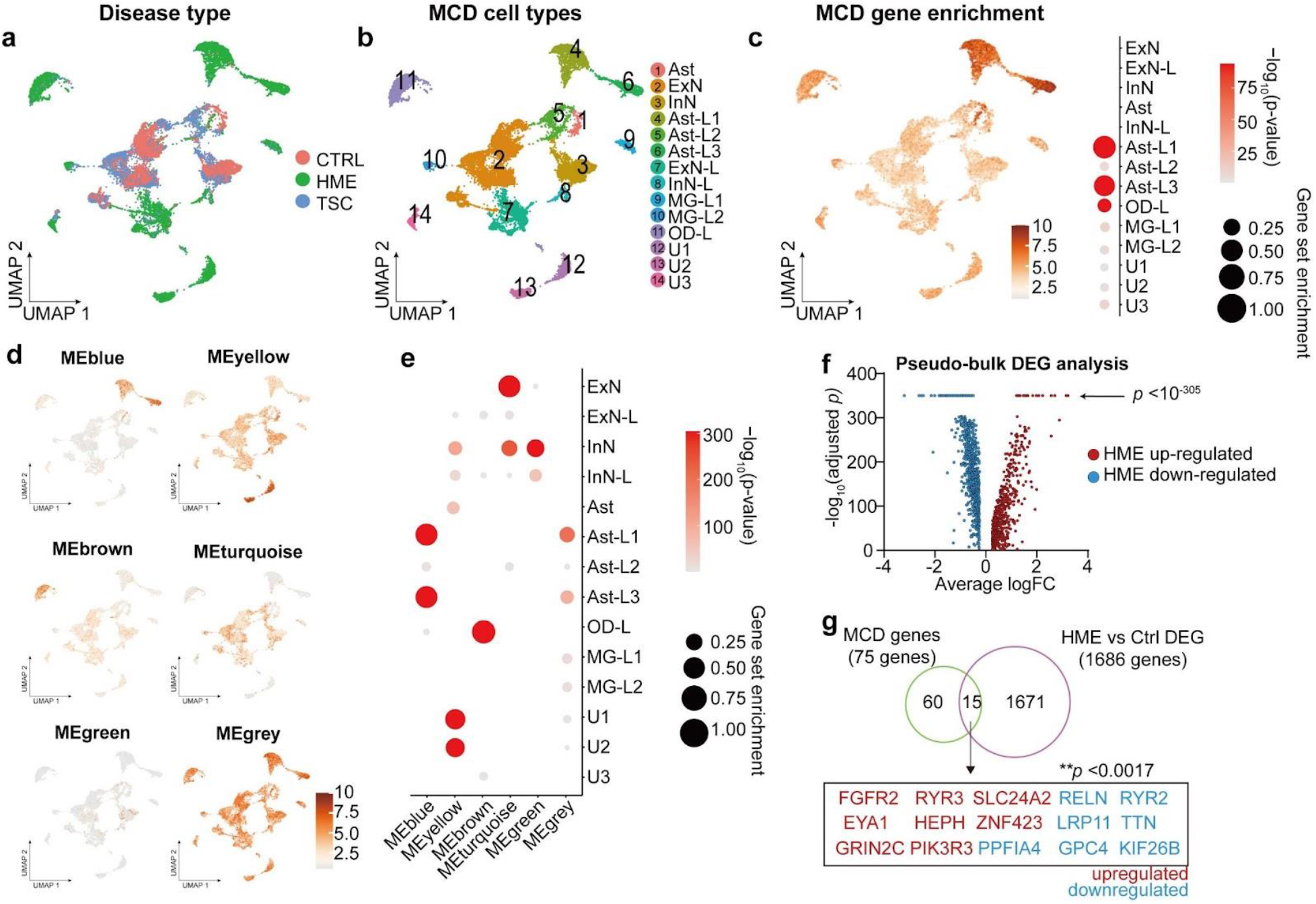
Single-nucleus transcriptomes showed MCD gene expression enriched in MCD-specific cell types. (a) UMAP for the single-nucleus transcriptome of 22067 nuclei from the cortical lesions of control (CTRL), hemimegalencephaly (HME), and tuberous sclerosis complex (TSC) brain. (b) Cell type classification. Ast: astrocyte, ExN: excitatory neuron, InN: inhibitory neuron, MG: microglia, OD: oligodendrocyte, U: unidentified. (c) The expression pattern of an eigengene made with all MCD genes and the quantification of enrichment based on cell types. (d) Six eigengenes decomposed from (c). (e) Quantification of the cell-type-specific enrichment in (d). (f) A volcano plot from DEG list of HME versus CTRL pseudo-bulk data. The genes having adjusted p < 10-305 were pointed by the arrow. (g) The MCD genes overlap with DEGs of HME in contrast to controls. A permutation test (10,000 times) shows a very rare chance (p < 0.0017) to show this overlap in a random sampling of 1686 genes from 19909 protein-coding genes used in these DEGs. Red or blue coloring of gene names indicates upregulated or downregulated DEGs in HMEs compared to CTRLs, respectively.

We noted that several of the HME clusters showed excessive expression of fibroblast growth factor receptor (FGFR) gene families, specifically *FGFR1* in cluster U1/2 in HME, *FGFR2/*3 in cluster Ast-L1/3 and OD-L, *EGFR* in Ast-L1/3 and U1/2, and *PDGFRA* in cluster U1/2 (Extended Data Fig. 5b,c). To identify the cell types expressing these genes, we performed RNA in situ hybridization in HME brain sections followed by hematoxylin-eosin staining. We found co-localization of these same *FGFR* family, *EGFR*, and *PDGFRA* transcripts with dysplastic cells (Extended Data Fig. 6). Previous experiments indicate that it is most often the dysplastic cells within HME and MCD that carry disease mutations^7^, suggesting an effect of these mutations on growth factor receptor expressions that correlates with dysplasia.

Next, we investigated the expression patterns of MCD genes in this HME/TSC snRNAseq dataset. An eigengene representing expression patterns of MCD genes was enriched in Ast-L1/3 and OD-L, which were labeled as dysplastic cells (Fig. 6c). Interestingly, the individual MCD genes displayed converging expression patterns resulting in six different eigengenes (Fig. 6d, gene members for each eigengene are described in Extended Data Fig. 7) which show distinct enrichment patterns across cell types (Fig. 6e), implying that membership of each eigengene may be associated with the pathophysiology of the corresponding dysplastic cell type in HMEs. We performed a pseudo-bulk DEG analysis comparing HME with CTRL and detected 590 up-regulated genes and 1096 down-regulated genes. Intriguingly, 20% (15/75) of MCD mutated genes in our list overlapped with DEGs of HME. Permutation testing suggests that this overlap is unlikely to have arisen by chance (Fig. 6f, see Methods). Taken together, many MCD genes are misregulated in MCD-specific cell types, suggesting that our MCD genes may play important roles in the pathogenesis of dysplastic cells in MCDs.

## Discussion

In this study, we use a multiomics approach to study the genetic landscape of MCD in the largest reported cohort to date. We confirmed the important role of mTOR/MAP kinase and glycosylation pathways, seen in about 60.5% of those with mutations. Moreover, our results also linked novel biological processes including gene expression, synaptic function, and calcium dynamics, which made up the other 39.5% of mutations. Nevertheless, only 76 of 317 patients showed one or more putative somatic mutations as a likely cause of MCD. There could be numerous causes for the relatively low solve rate in MCD, including the potential to miss very low VAF mutations and the contribution of complex mutations like structural variants or short tandem repeats polymorphism. Finally, although patients with environmental causes, syndromic, or inherited causes were excluded from our cohort, these factors could still contribute to MCD.

With our approach, we identified several recurrently-mutated genes not previously implicated in MCD. Confirming the remaining candidate and identifying further MCD candidate genes will require larger MCD cohorts. Including novel MCD candidate genes emerging from 300X WES into the 1000X Phase 3 AmpliSeq allowed both confirmation of mutations, a more accurate estimate of VAF, and identification of additional patients with these genes mutated that would have been perhaps missed with 300X WES. Functional validation by modeling mutations in embryonic mouse brains suggests that most candidate genes we identified are likely to contribute to disease. Perhaps it is not surprising that there are so many MCD genes, because such mutations may avoid embryonic lethality due to their expression in just a small subset of cells. Like with de novo germline mutations discovered in autism, we suggest that there could be dozens if not hundreds of additional MCD genes, based in part upon the low number of recurrently mutated genes ^42^.

The four gene networks, mTOR/MAP kinase, calcium dynamics, synapse, and gene expression, are intriguing, as they should play important roles for these genes both during brain development and homeostasis. All four pathways are critical both for corticogenesis during neurogenesis and neuronal migration, as well as neuronal excitability. For instance, calcium dynamics is shown to regulate cytoskeletal activity and excitability^43,44^. The genotypic information also showed correlations with clinical features, for instance, PET brain hypometabolism and abnormality in the neurological examination correlated with COSMIC DB variants, opening the possibility to predict genotype based on phenotype.

We also characterized the expression patterns of MCD genes in the developmentally normal and MCD brains at single-cell resolution. The cell types most strongly expressing candidate MCD genes include dorsal forebrain radial glial progenitors and their daughter excitatory neurons, as well as brain microglia, fitting well with the likely site of origin of somatic brain mutations^45^. Surprisingly, the dramatic gene dysregulation seen in the HME brain skewed the UMAP plots in ways that could not be accounted for simply by the VAF. The fact that the MCD genes also showed the strongest enrichment with these same clusters suggests that the MCD genes are very likely to have pivotal roles in the HME condition. Prior studies on MCD indicated that dysplastic cells express markers for both glia and neurons^46^. Our findings, however, suggest that MCD mutations drive critical roles predominantly in dividing radial glia, with profound effects on lineage and cellular dysplasia. To conclude, the MCD genes in patient brains found in our study demonstrated critical roles during cortical development, significantly correlate with patient phenotypes, and could open doors to novel treatments for MCDs.

## Online Methods

### Overview of the FCD cohort

This study is a multi-center international collaboration. We recruited a cohort of 317 individuals from the ‘FCD Neurogenetics Consortium’ (see the member list). These individuals were diagnosed with FCD type I, II, III, HME, or TSC and underwent surgical resection to treat drug-resistant epilepsy between 2013 and 2021. Any cases that underwent surgical resection due to environmental factors, for example, stroke, or acute trauma, were excluded. For each individual, resected brain tissue was collected, along with paired blood or saliva samples and parental samples, where available. Clinical history, pre- and post-operative brain imaging, histopathology, ILAE classification according to the surgical tissue pathology report, and Engel surgical outcome score (at least two years after surgery) were collected, when available.

### Informed consent and study approval

The study protocol was approved by the UC San Diego IRB (#140028). Informed consent was obtained from all participants or their legal guardians at the time of enrollment.

### DNA extraction

Pulverized cortical samples (∼0.3 g) were homogenized with a Pellet Pestle Motor (Kimble, #749540-0000) or Handheld Homogenizer Motor (Fisherbrand, #150) depending on the size of the tissue, and resuspended with 450 µL RLT buffer (Qiagen, #40724) in a 1.5 ml microcentrifuge tube (USA Scientific, #1615-5500). Homogenates were then vortexed for 1 minute and incubated at 70°C for 30 minutes. 50 μl Bond-Breaker TCEP solution (Thermo Scientific, #77720) and 120 mg stainless steel beads with 0.2 mm diameter (Next Advance, #SSB02) were added, and cellular disruption was performed for 5 minutes on a DisruptorGenie (Scientific industries). The supernatant was transferred to a DNA Mini Column from an AllPrep DNA/RNA Mini Kit (Qiagen, #80204) and centrifuged at 8500 xg for 30 seconds. The column was then washed with Buffer AW1 (kit-supplied), centrifuged at 8500 xg for 30 seconds and washed again with Buffer AW2 (kit-supplied), and then centrifuged at full speed for 2 minutes. The DNA was eluted two times with 50 μl of pre-heated (70°C) EB (kit-supplied) through centrifugation at 8,500 xg for 1 minute.

### MPAS and WES sequencing for somatic mutation candidates

Massive parallel amplicon sequencing (MPAS) and whole-exome sequencing (WES) were used at different phases to perform the genetic screening within available samples from the cohort. Customized AmpliSeq DNA panels for Illumina (Illumina, #20020495) were used for Massive Parallel Amplicon Sequencing^17^. 87 or 82 genes related to the mTOR pathway or curated based on the results of Phase 1 and 2, respectively, were subjected to the AmpliSeq design system; a list of designed genes is provided in Supplementary Table 2a-b. Two pools were designed for tiling the capture region. Genomic DNA from extracted tissue was diluted to 5 ng/uL in low TE provided in AmpliSeq Library PLUS (384 Reactions) kit (Illumina, #20019103). AmpliSeq was carried out following the manufacturer’s protocol (document #1000000036408v07). For amplification, 14 cycles each with 8 minutes were used. After amplification and FUPA treatment, libraries were barcoded with AmpliSeq CD Indexes (Illumina, #20031676) and pooled with similar molecular numbers based on measurements made with a Qubit dsDNA High Sensitivity kit (Thermo Fisher Scientific, #Q32854) and a plate reader (Eppendorf, PlateReader AF2200). The pooled libraries were subjected to Illumina NovaSeq 6000 platform for PE150 sequencing. The AmpliSeq design in the ‘Phase 1’ is under the design ID IAA7610, and the AmpliSeq design in ‘Phase 3’ is under the design ID IAA26010.

Genomic DNA (∼ 1.0 μg) was prepared for whole-exome sequencing, and libraries were captured using the Agilent SureSelect XT Human All Exon v.5 or Nextera DNA Exome kits. Then, 100, 125, or 150 bp paired-end reads (median insert size ∼ 210 bp) were generated using the Illumina HiSeq X 2500 platform. The sequencing experiments were designed to yield three datasets of ∼ 100X coverage on each sample, with a coverage goal of 300X from the brain and 100X from blood/saliva.

### Somatic variant calling from MPAS and WES

Reads were aligned to GRCh37 using BWA (version 3.7.16a), sorted per each read group, and merged into a single BAM file with Sambamba (version 0.6.7). The merged BAM files were marked for duplicate reads using PICARD (v2.12.1), duplicated reads were not removed for MPAS because of the nature of the method. Then, we performed indel realignment and base quality recalibration using GATK (v3.7–0), resulting in the final uniformed processed BAM files.

Both tissue-specific and tissue-shared mosaic variants were called from the MPAS and WES sequencing data. MPAS and WES variants were called according to the availability of the control tissue. Brain- and blood/saliva-specific variants were called using MuTect2 (GATK3.8) paired mode and Strelka2 somatic mode^47^; the BAM files from the brain sample (combined and non-combined from independent sequencing libraries) and blood/saliva samples were treated as “tumor-normal” and “normal-tumor” pairs separately and cross-compared between each other. Variants called by both callers were listed. Mosaic variants shared between the brain and fibroblast samples were called using the single mode of MosaicHunter^11^ by either combining all brain replicates or calling each separate sample. Variants that passed all the MosaicHunter filters also were listed. Somatic variants from WES data were further called by GATK (v3.7–0) haplotypecaller with ploidy parameter set to 50, followed by a series of heuristic filters described as the best-practice by the Brain somatic mosaicism network^12^, tissue-shared variants were called by the combination of MuTect2^48^ (GATK 3.8) single-mode and DeepMosaic^10^.

A union of different pipelines was selected to get maximum sensitivity. Mosaic candidates from the combined lists were further filtered using the following criteria: (i) the variant had more than 3 reads for the alternative allele; (ii) the variant was not present in UCSC repeat masker or segmental duplications; (iii) the variant was at least 2 bp away from a homopolymeric tract; and (iv) the variant exhibited a gnomAD allele frequency lower than 0.001. Variants that exist in the 1000 genome project (phase 3) also were excluded from the analysis. Variants from both exome data sources were tested and a combination of tissue-specific mosaic variants and tissue-shared mosaic variants were collected and the credible interval of VAFs was calculated using a Bayesian-based method described previously^49^. To filter for candidate disease-causing variants for FCD, we further filtered out synonymous variants in coding regions, variants with CADD Phred score < 25, and candidates that fell out of coding regions and were not predicted to affect splicing by ANNOVAR.

### False discovery estimation

To calculate the false discovery of random variants detected in normal samples, we incorporated 75 normal control samples (71 brains and 4 other organs) previously sequenced with 250-300X WGS, which should provide similar sensitivity as our exomes, the deep WGS were generated by efforts from the NIMH Brain Somatic Mosaicism Consortium^12^, from controls^17^, and from our recent mutation detection pipeline^18^. Variants were filtered based on the identical criteria as described in the above data analysis part, with >0.01 VAF, all on exonic regions defined by NCBI, and CADD score >25. While 13 variants remain positive from this pipeline from the 75 samples (0.17 per control), 306 candidate variants were determined in our 134 MCD exomes (2.28 per MCD case), which lead to an estimated 7.59% per sample false discovery rate (Supplementary Table 5).

### Orthogonal validation and quantification of mosaic mutations with targeted amplicon sequencing

Targeted amplicon sequencing (TASeq) with Illumina TruSeq was performed with a coverage goal of >1000X for 554 candidate variants detected by computational pipelines described above for both MPAS and WES, to experimentally validate the mosaic candidates before functional assessment. PCR products for sequencing were designed with a target length of 160-190 bp with primers being at least 60 bp away from the base of interest. Primers were designed using the command-line tool of Primer3^50,51^ with a Python (v3.7.3) wrapper^13,14^. PCR was performed according to standard procedures using GoTaq Colorless Master Mix (Promega, M7832) on sperm, blood, and an unrelated control. Amplicons were enzymatically cleaned with ExoI (NEB, M0293S) and SAP (NEB, M0371S) treatment. Following normalization with the Qubit HS Kit (ThermFisher Scientific, Q33231), amplification products were processed according to the manufacturer’s protocol with AMPure XP beads (Beckman Coulter, A63882) at a ratio of 1.2x. Library preparation was performed according to the manufacturer’s protocol using a Kapa Hyper Prep Kit (Kapa Biosystems, KK8501) and barcoded independently with unique dual indexes (IDT for Illumina, 20022370). The libraries were sequenced on Illumina HiSeq 4000 or NovaSeq 6000 platform with 100 bp paired-end reads.

### Mutational signature analysis

Mutational signature analysis was performed using a web-based somatic mutation analysis toolkit (Mutalisk)^52^. PCAWG SigProfiler full screening model was used.

### STRING analysis

STRING analysis was performed by STRING v11^22^. A total of 75 MCD genes were loaded as input and MCL clustering was performed. The terms in Gene Ontology (GO), KEGG pathways, and Top 10 terms GO or KEGG pathways were shown in Fig. 2b. If there are less than 10 terms for those terms (such as clusters 3 and 4 in Fig. 2), we included all the terms in GO or KEGG pathways, Local network cluster (STRING), Reactome pathways, and Disease-gene associations (DISEASES) to show the enriched terms. Visualization was performed by Cytoscape v3.9.

### ClueGO analysis

Visualization of the functionally grouped biological terms was performed by ClueGO v2.5 ^53^, a Cytoscape plug-in. A total of 75 MCD genes from Fig. 2 were loaded and GO terms in the ‘Biological Process’ category were used for visualization. Terms with a p < 0.01, a minimum count of 3, and an enrichment factor > 1.5, are grouped into clusters based on membership similarities.

### Animals

Pregnant Crl: CD1(ICR) mice for mouse modeling were purchased from Charles River Laboratory. All mice used were maintained under standard group housing laboratory conditions with 12 hours light/dark cycle and free access to food and water. The age and number of mice used for each experiment are detailed in the figure legends. The sex of the embryos used was not tested. All work with mice was performed in accordance with UCSD IACUC protocol S15113.

### DNA constructs

*RRAGA, KLHL22*, and *RHOA* ORF regions were amplified from the hORFeome library and inserted into the pCIG2 (pCAG-IRES-GFP) vector. *GRIN2C* ORF region was purchased from DNASU Plasmid Repository in Arizona State University Biodesign Institute. All sequences of clones were confirmed by sanger sequencing.

### In utero electroporation

In utero electroporation was performed as described previously^54^ with modifications as follows. Endotoxin-free plasmids (0.5–1 μg) plus 0.1% Fast Green (Sigma, catalog no. 7252) was injected into one lateral ventricle of E14.5 embryos. Electroporation was performed by placing the anode on the side of the DNA injection and the cathode on the other side of the head to target cortical progenitors. Four pulses of 45 V for 50 ms with 455-ms intervals were used.

### Mouse brain section preparation

An E18 mouse brain is fixed in 4% paraformaldehyde (PFA) for 2 hrs. For the P21 mouse brain, a mouse was anesthetized by isoflurane and perfused by cold 1X PBS for 8 min and following 4% cold PFA for 8 min. The brains were dehydrated in 30% sucrose in 1x PBS for 48 hrs and embedded in Tissue-Tek optimal cutting temperature compound and frozen on dry ice. A frozen block was sectioned with 20 um thickness in a cryostat (CryoStar NX70, Thermo Fisher Scientific) and placed on sliding glass. The attached sections were dried on a 50 °C heating block for 3 hrs.

### Immunofluorescence staining and imaging

A section was rehydrated and washed by 1X PBS for 10 min 3 times, permeabilized in PBST (0.3% Triton X-100 in 1X PBS) for 10 min, and blocked by blocking solution (5% normal BSA in 1X PBS) for 2 hrs in room temperature. Sections were stained with diluted primary antibodies in the blocking solution overnight at 4 °C. The next day, the sections were washed with PBST for 5 min three times and stained with secondary antibodies in blocking solution for 2 hrs in RT. Blocking solution was dropped off from the slides and nuclei staining with DAPI solution (0.1ug/ml of DAPI in PBST) was performed for 15 min. The slides were mounted with DAKO fluorescent mount solution (catalog no. S3023). Zeiss 880 Airyscan Confocal is used for imaging according to the manufacturer’s instructions.

### Antibodies

phospho-S6 (1:800 dilution, catalog no. 5364S ;Cell Signaling, AB_10694233), NeuN (1:100, MAB377X; Sigma-Aldrich, AB_2149209), GFP (1:500, catalog no. GFP-1020, Aves Labs, AB_10000240), Alexa Fluor Goat 488 chicken IgY (H+L) (1:1,000 dilution, catalog no. A-11039, AB_2534096), Alexa Fluor 594 donkey anti-rabbit lgG (H+L) (1:1,000, catalog no. R37119, AB_2556547).

### Genotype-phenotype association

The functional modules to be tested were selected based on the enriched GO terms (Fig. 2 and Extended Data Fig. 4). A given candidate MCD gene was assigned as a member to one or multiple modules based on GO terms related to the given gene (results summarized in Supplementary Table 3c). Subsequently, a given patient became a member of one (or multiple) functional module(s) if the genes detected in that patient were assigned to that (those) functional module(s). All available clinical information on the patient was collected and harmonized using ILAE terms (summarized in Supplementary Table 4). Pearson correlation coefficients were calculated by cor.test() function in R. The value of correlation coefficients were displayed as colors in the heatmap of Fig. 4. If two groups with binary values were used for calculation, Phi coefficient was used.

### Single-nucleus RNA sequencing

A fresh-frozen brain tissue (∼50 mg) was placed into a glass dounce homogenizer containing 1 ml cold lysis buffer (0.05 % (v/v) NP-40, 10 mM Tris (pH 7.4), 3 mM MgCl_2_, 10 mM NaCl) and dounce 10 times with a loose pestle and following 10 times with a tight pestle. The homogenate was incubated for 10 min in RT. 9 ml of wash buffer (1% BSA in 1X PBS) was added to the homogenate and filtered by a 30 um cell strainer. The strained homogenate was spun down in 500 g to remove the supernatant. The pellet was resuspended by 5 ml of wash buffer. Straining, spinning down steps was performed once more, and the pellet was resuspended into 500 ul of wash buffer. 10 ul of nuclei resuspension was mixed with counting solution (0.02 % Tween 20, 0.1ug/ml DAPI, 1% BSA in 1X PBS) and nuclei density was measured by manual nuclei counting using DAPI signal. The resuspension was diluted by wash buffer to make the desired concentration (800∼1000 nuclei/ul). 1∼4 samples were pooled together targeting 10000 nuclei per reaction. Gel beads emulsion (GEM) generation, cDNA, and sequencing library constructions were performed in accordance with instructions in the Chromium Single Cell 3’ Reagent Kits User Guide (v3.1). A library pool was sequenced with 800 million read pairs using NovaSeq 6000.

### Single-nucleus RNAseq bioinformatics pipeline

Fastq files from single-nucleus libraries were processed through Cell Ranger (v6.0.2) analysis pipeline with –include-introns option and hg19 reference genome. Pooled library was demultiplexed and singlets were taken by demuxlet. Seurat (v4) package was used to handle single nuclei data objects. Protein coding genes were used for further downstream analysis. Nuclei passed a control filter (number of genes > 500, number of reads >1000, percentage of mitochondrial gene < 10%) was used for downstream analysis. Data were normalized and scaled with the most variable 5000 features using the ‘NormalizeData’ and ‘ScaleData’ functions. Dimensionality reduction by PCA and UMAP embedding was performed using runPCA and runUMAP function. Clustering was performed by FindNeighbors and FindClusters function. Cell type identification was performed using known cell type markers expressed in the brain including excitatory/inhibitory neuron, astrocyte, oligodendrocyte, microglia, and endothelial cell markers as well as using positive markers found by FindAllMarkers function with 3000 most variable features in scaled data.

### Weighted gene co-expression network analysis

‘r-wgcna’ package (v1.69) was used for WGCNA according to instructions (PMID: 19114008). Briefly, a similarity matrix was generated based on Pearson’s correlation coefficient value among the top 3000 variable features in single-nucleus transcriptome data, which was used to calculate the subsequently signed type of network adjacency matrix. Next, the topological overlap matrix (TOM) and the corresponding dissimilarity (1-TOM) value were generated from the adjacency matrix. Finally gene modules were generated by ‘cutreeDynamic’ function with ‘tree’ method, minAbsSplitHeight = 0.9 and minClusterSize = 30 option. Similar gene modules were merged by ‘mergeCloseModules’ function with cutHeight = 0.25. String analysis was performed using each gene module for the identification of the given module’s functional characteristics.

### RNAscope

We used published methods and purchased target probes for genes of interest containing an 18-25 base region complementary to the target, as spacer sequencing, and a 14 base Z-tail sequence^55^, including RNA pol III positive control and random sequence negative control, following the manufacturer recommendations (Advanced Cell Diagnostics, Hayward, CA). Images were acquired on a Leica STED Sp8 with Falcon microscope.

### Permutation analysis for the enrichment of MCD genes

To test the enrichment of differentially expressed MCD genes in RNA sequencing against a random distribution, we designed a permutation analysis. All human genes used in the single-cell RNA-seq analysis (n=19909) were randomly shuffled 10,000 times and the same number of genes as described in the differential expression analysis (n=1686) was selected for each shuffle. The number of overlaps between each shuffle and the MCD candidates was compared and the number of overlaps was used as the outcome and a null distribution was generated from the 10,000 shuffles. All 75 positively validated MCD genes are confirmed to be existing in the initial gene list. After 10,000 permutations, the permutation p-value was calculated with numbers >= observed overlap (p=0.0017 for the data shown in the main text).

### Statistical analyses

Statistical analyses were performed by R or Prism 8 (GraphPad Software). Two-way ANOVA and Sidak multiple comparisons were performed in Fig 3b with p-values of interaction between genotype and bin factor. \*\*\*\**p* < 0.0001, \*\*\**p* < 0.001, \*\**p* < 0.01, \**p* < 0.05.

## Code and data availability

Code to generate the figures and analyze the data are publically available on GitHub (https://github.com/shishenyxx/MCD_mosaic). WES and AmpliSeq data are deployed on NIMH Data Archive under study number 1484 “Comprehensive multiomic profiling of somatic mutations in malformations of cortical development” and SRA under accession number PRJNA821916: “Comprehensive multiomic profiling of somatic mutations in malformations of cortical development”. The snRNAseq R object was deposited in Single Cell Portal (https://singlecell.broadinstitute.org/single_cell/study/SCP1815/comprehensive-multiomic-profiling-of-somatic-mutations-in-malformations-of-cortical-development#study-download).

## Acknowledgments

AmpliSeq, TASeq, and snRNAseq were supported by NIH P30CA023100 and S10OD026929 at the UCSD IGM Genomics Center. Rady Children’s Institute for Genomic Medicine, Broad Institute (U54HG003067, UM1HG008900), the Yale Center for Mendelian Disorders (U54HG006504), and the New York Genome Center provided whole-exome sequencing. UCSD Microscopy core (NINDS P30NS047101) provided imaging support. CC was supported by a 2021 Brain & Behavior Research Foundation Young Investigator Grant. This study was supported by the NIH (NIMH U01MH108898 and R01MH124890 to JGG and GWM, and NIA R21AG070462, NINDS R01NS083823 to JGG). We thank Stéphanie Baulac and Sara Baldassari for sharing unpublished exome data.

## Author contributions

C.C., X.Y., and J.G.G. designed the study. C.C., S.M., and S.K. conducted functional validation. C.B., V.S., A.N., E.R., C.C., and G.H. coordinated the clinical database. X.Y., C.C., M.W.B., L.L.B., R.D.G., J.G., M.X., A.P.L.M., and K.N.J. organized, handled, and sequenced human samples. X.Y., C.C., T.B., X.X., and B.C. performed bioinformatics and data analysis. C.C. and K.I.V. performed the RNAscope experiment. C.D., H.W.P., C.A.B.G., S.H.K., H.K., A.S., C.A.H., C.G., C.A.G., S.S., M.N., D.D.G., K.I., Y.T., R.C., J.T., V.C., R.G., O.D., W.A.S., H.R.M., and G.W.M. provided resected brain tissues and clinical data from FCD patients. C.C., X.Y., and J.G.G. wrote the manuscript. All authors read and commented on the manuscript before submission.

## Supplementary Table Descriptions

**Supplementary Table 1. The cohort list and corresponding sequencing methods**. The 327 cases are listed in each row and corresponding sequencing methods used for a given sample were described.

**Supplementary Table 2. AmpliSeq primer pool designs** (a) Ampliseq primer pool design used in phase 1. (b) Ampliseq primer pool design used in phase 3.

**Supplementary Table 3. The summary of SNV calls across the three phases of genetic discovery**. (a) 1811 raw calls derived from the combination of variant callers described in Extended Data Fig. 1. (b) 554 input SNV calls participated in TASeq quantification. (c) Validated brain somatic SNV calls from (b). (d) Annotation table of the genes listed in (c) based on GO terms.

**Supplementary Table 4. The summary of phenotype and genotype information for the ‘genetically solved’ cases**.

**Supplementary Table 5. The summary table used for false discovery estimation**.

**Extended Data Fig. 1.**
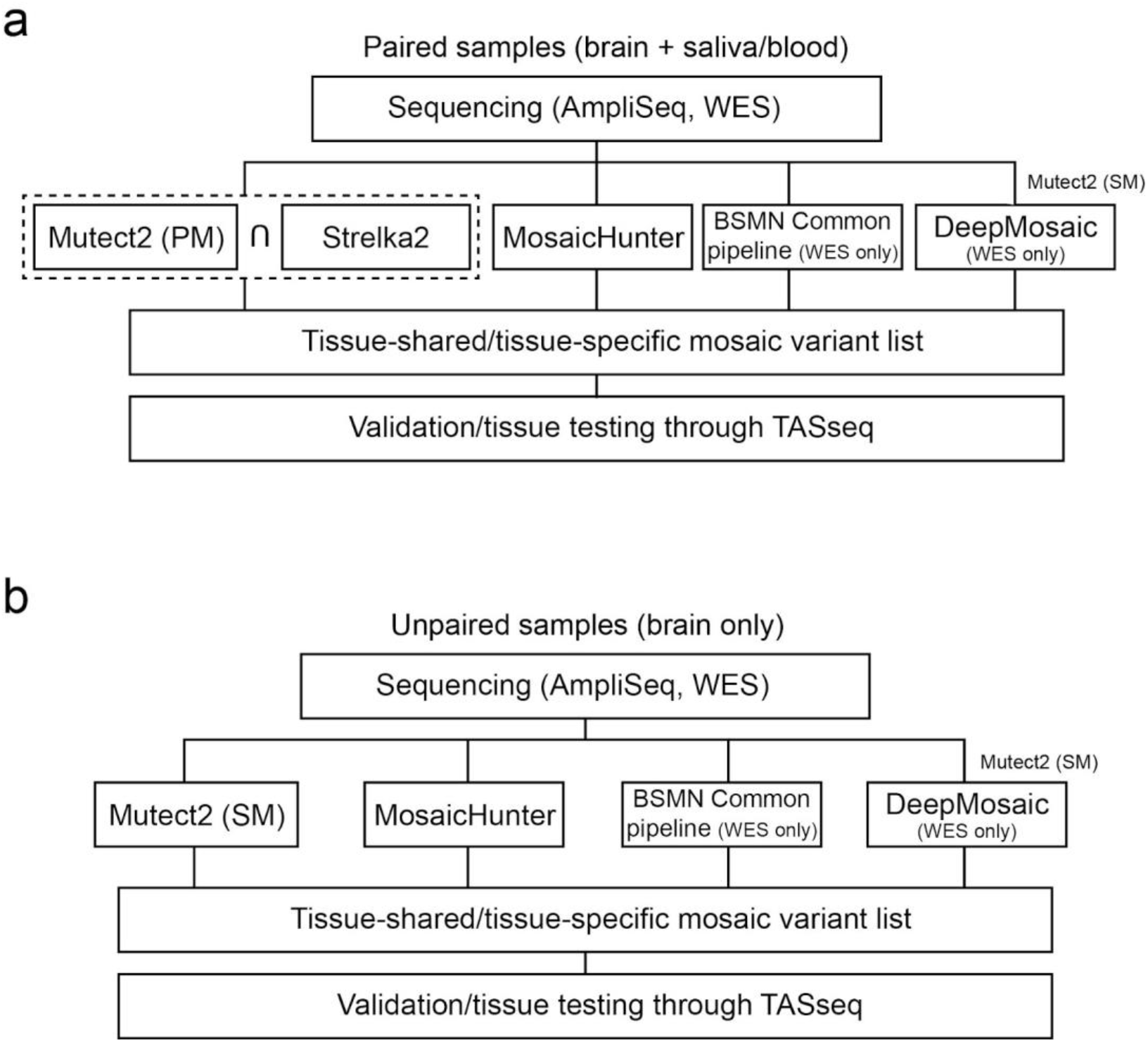
Bioinformatic pipeline to detect somatic SNVs in the MCD cohort. (a) The pipeline for paired samples. Notably, the dashed square indicates that the sharing variants between MuTect2 paired mode and Strelka2 were used for the downstream analysis. BSMN common pipeline and DeepMosaic were used only for WES datasets. The DeepMosaic input variants were generated by MuTect2 single mode. (b) The pipeline for unpaired samples. The pipeline is similar except that MuTect2 single mode without Strelka2 is used. PM: paired mode, SM: single mode.

**Extended Data Fig. 2.**
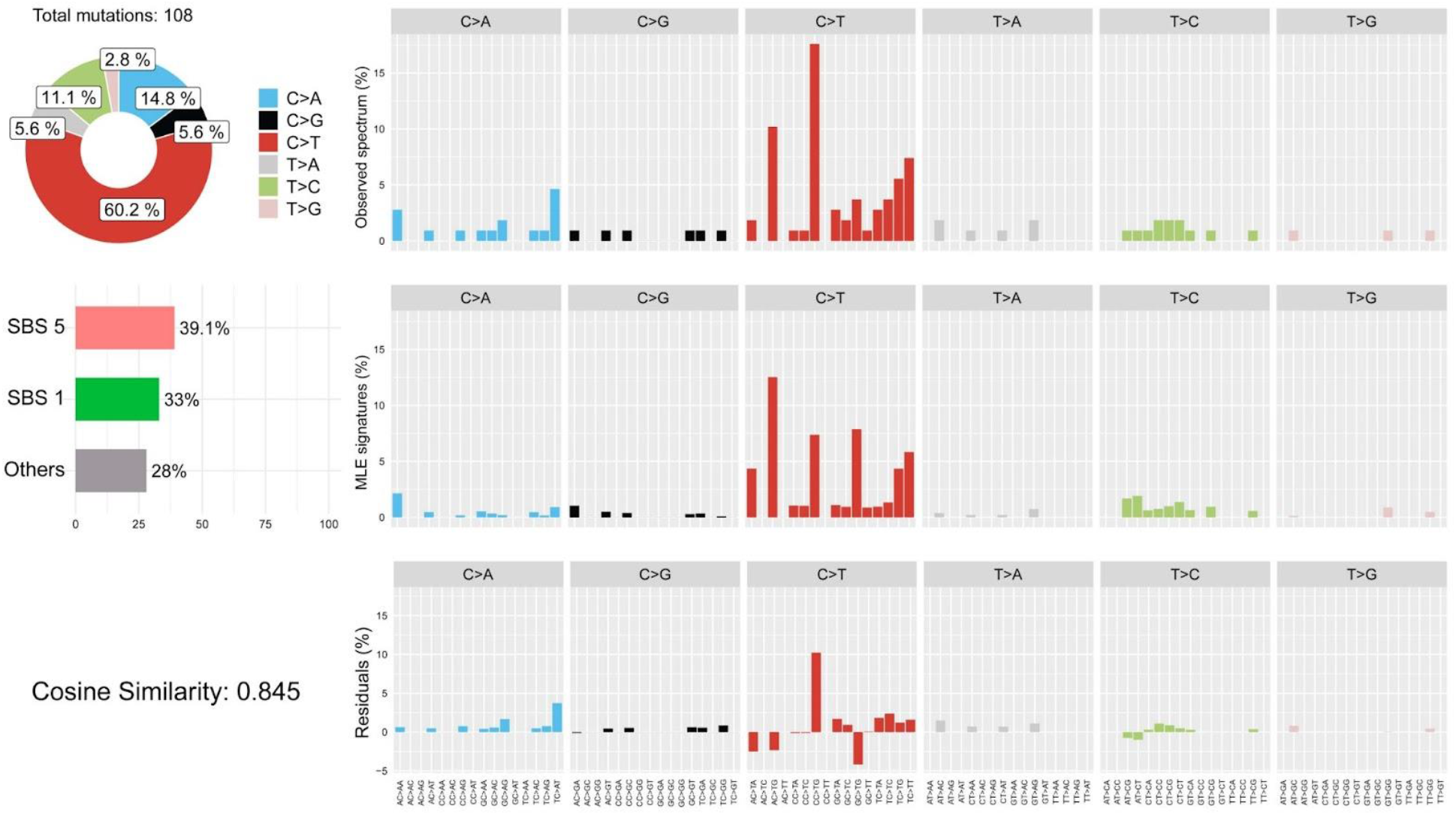
Mutational signature analysis through Mutalisk shows cell-division-related clock-like signatures in the MCD cohort. SBS5 (39.1%) and SBS1 (33%) are clock-like mutational signatures. SBS1 especially correlates with cell division and mitosis of stem cells.

**Extended Data Fig. 3.**
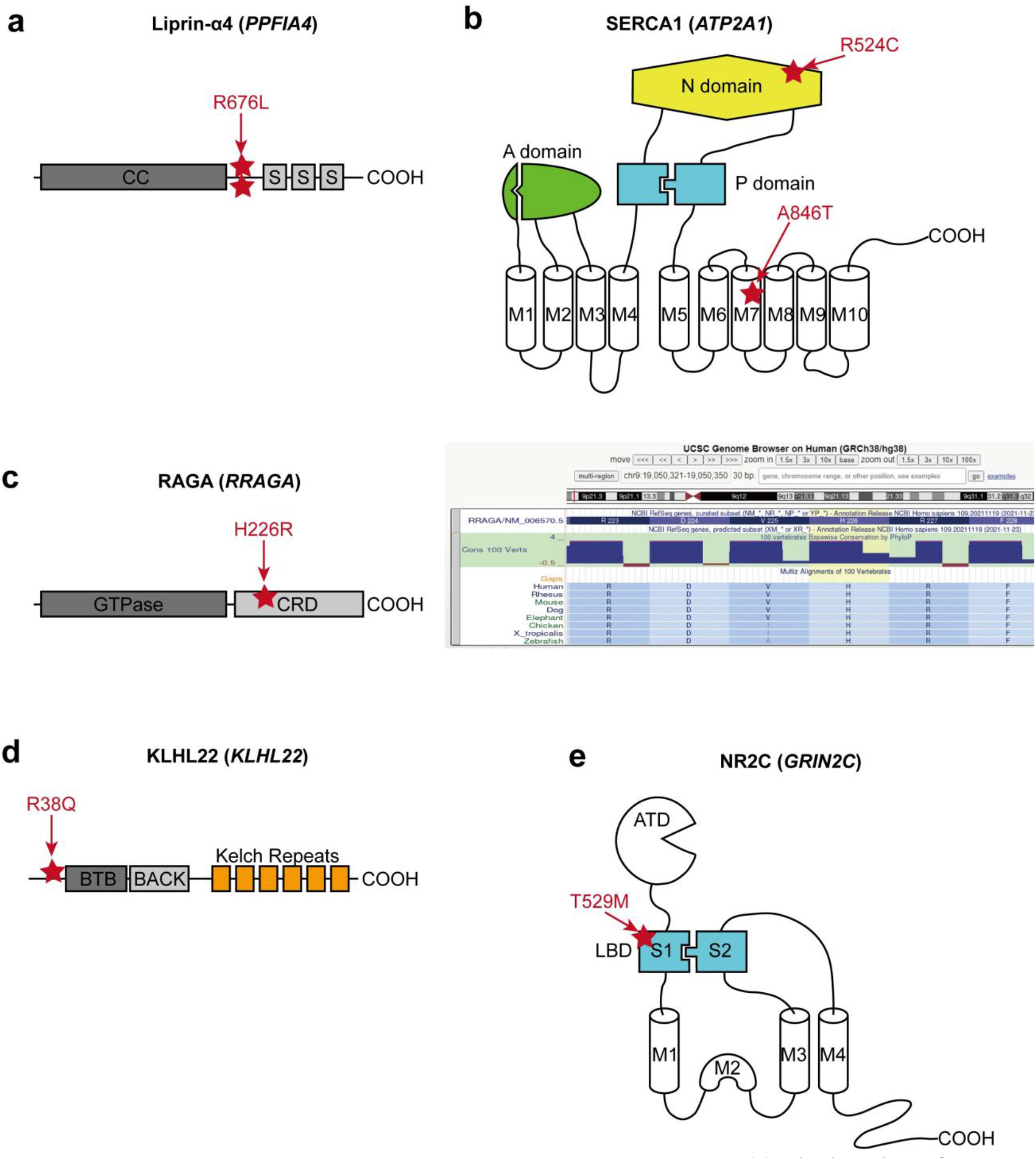
The locations of the selected MCD variants. (a) The location of two recurrent variant calls is at the same position between the coiled-coil domain (CC) and the first SAM domain (S) of Liprin-α4. (b) Two different variants in SERCA1. p.R524C mutation is at the nucleotide ATP-binding (N) domain, whereas the pA846T variant is in the 7th transmembrane (M7) domain. A: Actuator domain, P: Phosphorylation domain, M: Transmembrane domain. (c) Left: The location of p.H226R variant in RAGA protein. GTPase: GTPase domain, CRD: C-terminal roadblock domain. Right: UCSC genome browser screenshot describing that p.H226 is a conserved site across all vertebrates. (d) The location of p.R38Q variant in the N-terminal region before BTB (Broad-Complex, Tramtrack, and Bric-à-brac) domain of KLHL22. (e) A variant in the S1 domain of NR2C. S1 and S2 together make the ligand-binding domain (LBD), the target of glutamate. ATD: Amino terminal domain.

**Extended Data Fig. 4.**
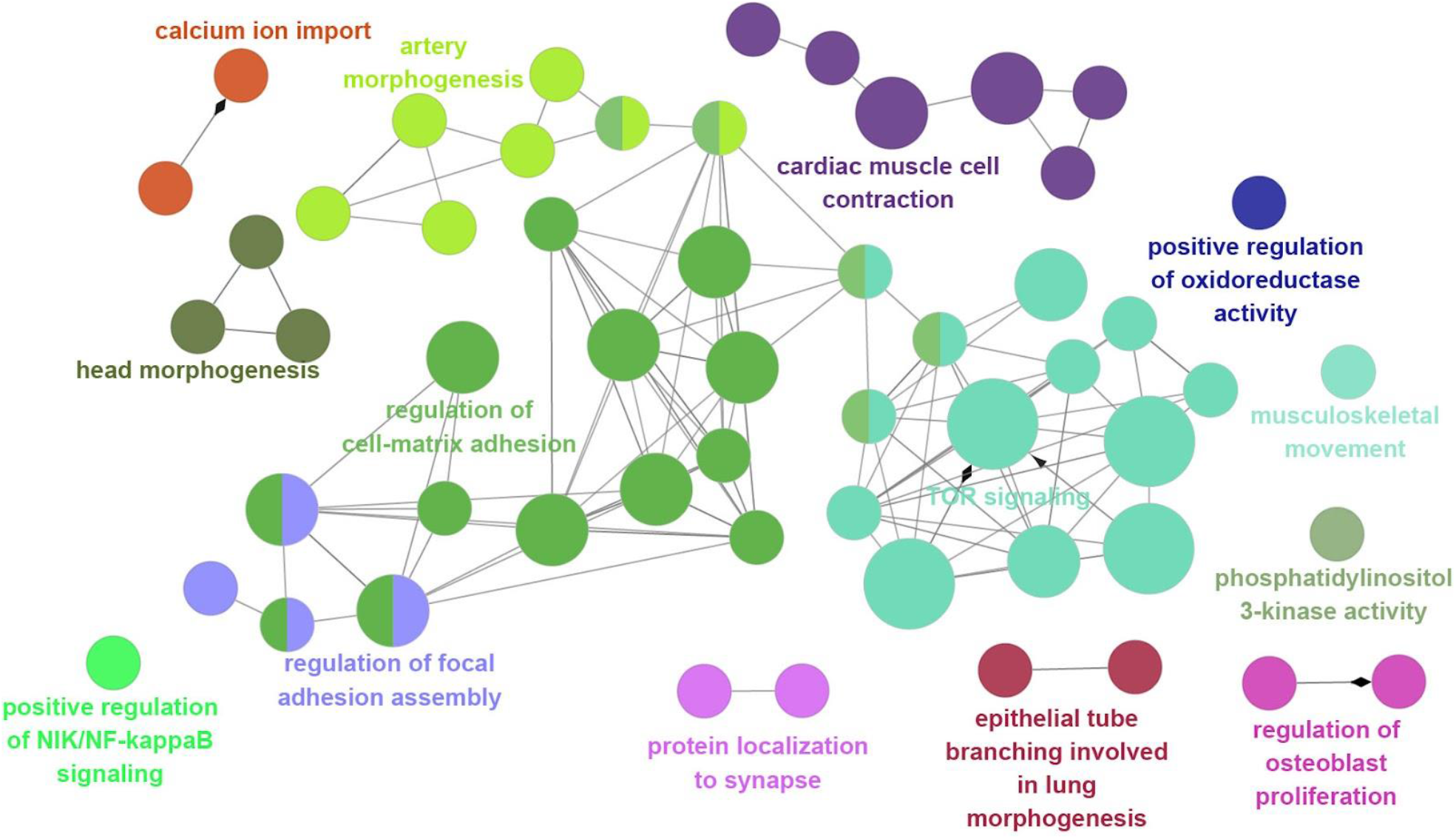
The ClueGO analysis using the MCD genes result identifies the biological processes and molecular pathways. The main cluster is related to TOR signaling, regulation of cell-matrix adhesion, regulation of focal adhesion assembly, and artery morphogenesis. Notably, there are also isolated clusters that were not covered in previous studies, for example, cardiac muscle cell contraction, calcium ion import, and protein localization to the synapse. Term p-value with Bonferroni correction was reflected in node size (Large: *p* < 0.0005, medium: *p* < 0.05, small: *p* < 0.1).

**Extended Data Fig. 5.**
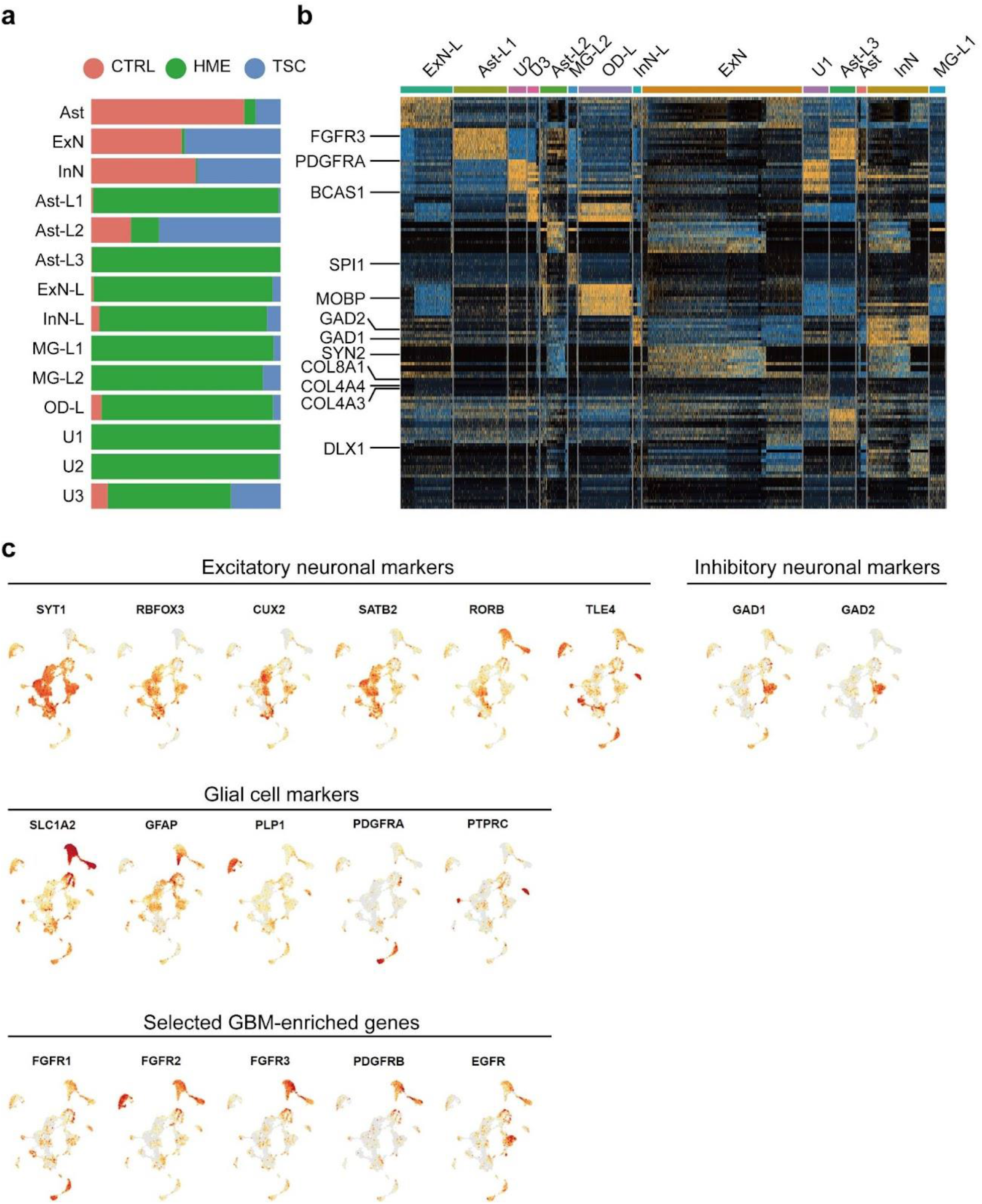
Cell-type identification by DEGs and known marker gene expression in the MCD snRNAseq dataset. (a) MCD prefix was used for the clusters that have less than 25 % of control origin. (b) DEG analysis using FindAllMarker function in Seurat v4 package. The top 10 genes for each cluster were presented. Several notable genes helping to define major cell types were labeled on the left side. Note that *FGFR3* and *PDGFRA* are up-regulated in Ast-L1/3 and U1/2/3, respectively, implying that these genes can be the markers for MCD-dominant clusters. (c) Selected markers for major cell types in the human cortex. *CUX1, CUX2* for upper layer excitatory neuronal markers, SATB2 for layer 4 excitatory neuronal marker, *RORB, FEZF2, BCL11B, FOXP2, ROBO2* for deep layer-specific markers, *GAD1, GAD2, DLX6, RELN* for inhibitory neuronal markers, *GFAP, SLC1A2, SLC1A3, MMD2* for astrocyte markers, *PTPRC* for the microglial marker, *OLIG1, OLIG2, MOBP, PLP1* for oligodendrocyte markers, *FGFR1, FGFR2, FGFR3, PDGFRB, EGFR* for the selected GBM-enriched genes covering subsets of MCD-enriched clusters.

**Extended Data Fig. 6.**
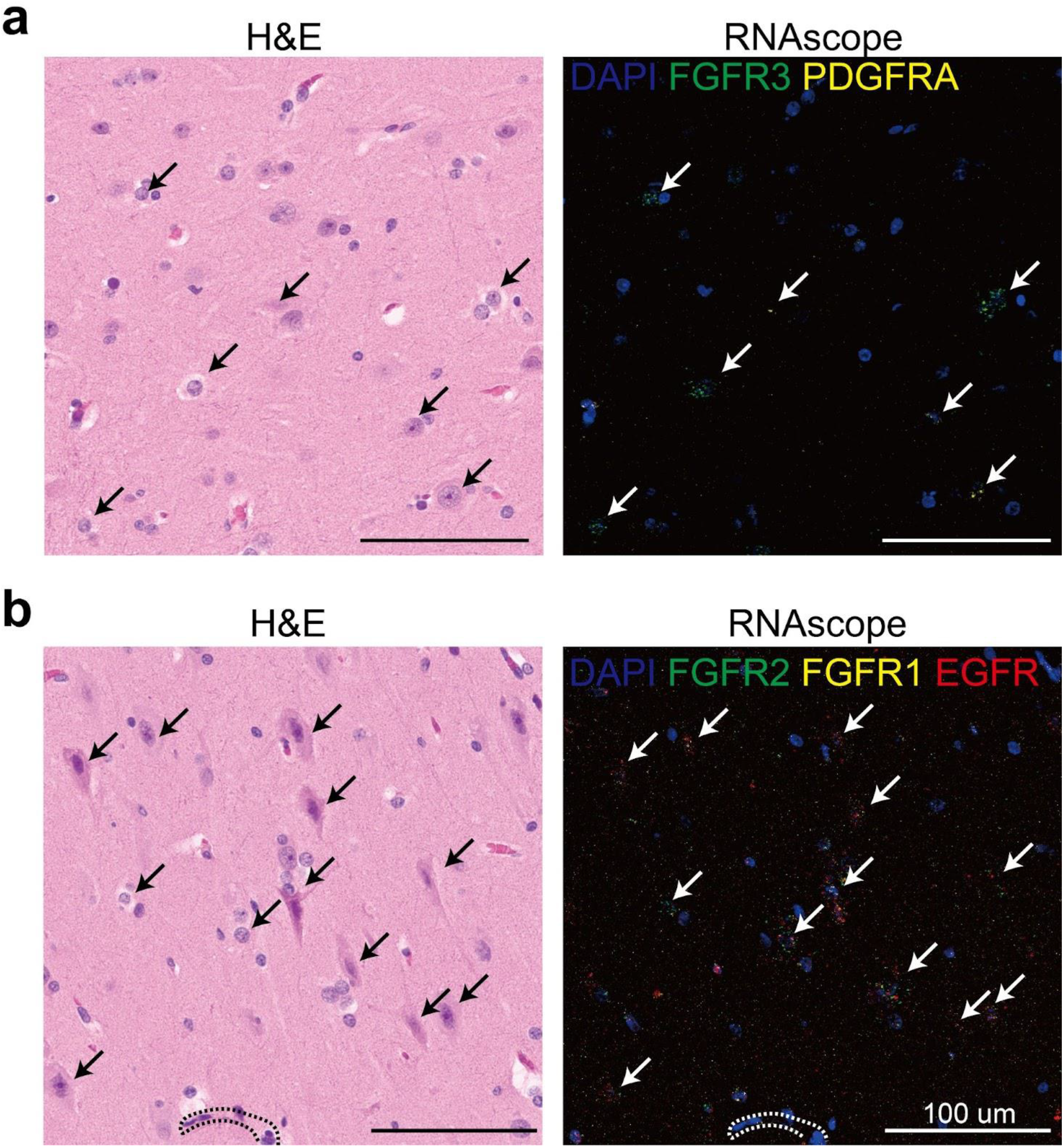
The validation of the snRNAseq result from HME-6593 shows MCD dominant clusters are highly correlated with dysplastic cells in MCD. (a) H&E staining (left) and RNAscope (right) staining results in several MCD-dominant markers (*FGFR2, FGFR1, EGFR*) in the same formaldehyde-embedded-paraffin-fixed section. (b) H&E and RNAscope result in another section with different RNA probes (*FGFR3* and *PDGFRA*) enriched in MCD clusters. Dashed lines indicate blood vessels. White/black arrows are pointing to the dysplastic cells.

**Extended Data Fig. 7.**
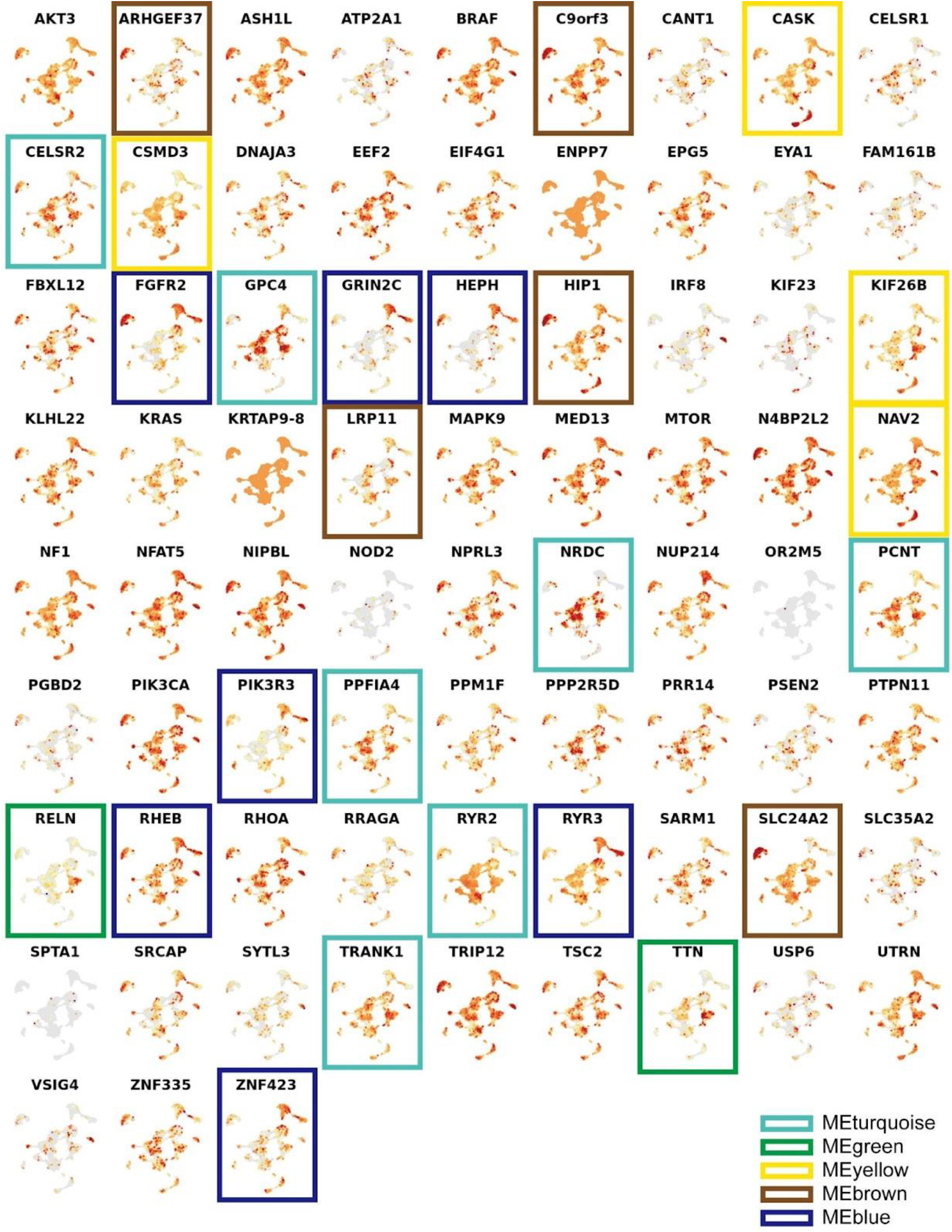
Expression patterns of individual MCD genes in the MCD snRNAseq dataset. The gene members of each eigen module shown in Fig. 6d were colored according to the name of a given eigengene.

## Focal Cortical Dysplasia Neurogenetics Consortium (Additional Members)

Dr. Yasemin Alanay, Division of Pediatric Genetics, Acibadem Hospital, Istanbul, Turkey Dr. Seema Kapoor, Division of Genetics, Genetic & Metabolic Lab, Lok Nayak Hospital & Maualana Azad Medical Center, Pakistan

Dr. Georgia Ramantani, Dr. Thomas Feuerstein, Albert-Ludwigs University, Freiburg, Germany Dr. Ingmar Blumcke, Dr. Robyn Busch, Dr. Zhong Ying, Department of Neuropathology, University Hopsital Erlangen, Germany

Dr. Vadym Biloshytsky, Dr. Kostiantyn Kostiuk, Dr. Eugene Pedachenko, A. Romodanov Institute of Neurosurgery, Kyiv, Ukraine

Dr. Marilyn Jones, Diane Masser-Frye, Rady Children’s Hospital, San Diego, CA

Dr. Ingo Helbig, Dr. Benjamin C. Kennedy, Division of Neurology, Children’s Hospital Philadelphia, PA

Dr. Judy Liu, Dr. Felix Chan, Department of Molecular Biology, ell Biology, and Biochemistry, Department of Neurology, Brown University, RI

Dr. Darcy Krueger, Department of Clinical Pediatrics and Neurology, Cincinnati Children’s Hospital, OH

Dr. Richard Frye, Dr. Angus Wilfong, Dr. David Adelson, Barrow Neurological Institute at Phoenix Children’s Hospital, U Arizona College of Medicine, Phoenix, AZ

Dr. William Gaillard, Dr. Chima Oluigbo, Children’s National Hospital, Washington DC

Dr. Anne Anderson, Dept of Pediatrics, Baylor College of Medicine, Texas Children’s Hospital,

Houston, TX

“,

## Brain Somatic Mosaicism Network

Boston Children’s Hospital: Alice Lee, August Yue Huang, Alissa D’Gama, Caroline Dias, Christopher A. Walsh, Eduardo Maury, Javier Ganz, Michael Lodato, Michael Miller, Pengpeng Li, Rachel Rodin, Rebeca Borges-Monroy, Robert Hill, Sara Bizzotto, Sattar Khoshkhoo, Sonia Kim, Zinan Zhou

Harvard University: Alice Lee, Alison Barton, Alon Galor, Chong Chu, Craig Bohrson, Doga Gulhan, Eduardo Maury, Elaine Lim, Euncheon Lim, Giorgio Melloni, Isidro Cortes, Jake Lee, Joe Luquette, Lixing Yang, Maxwell Sherman, Michael Coulter, Minseok Kwon, Peter J. Park, Rebeca Borges-Monroy, Semin Lee, Sonia Kim, Soo Lee, Vinary Viswanadham, Yanmei Dou

Icahn School of Medicine at Mt. Sinai: Andrew J. Chess, Attila Jones, Chaggai Rosenbluh, Schahram Akbarian

Kennedy Krieger Institute: Ben Langmead, Jeremy Thorpe, Sean Cho

Lieber Institute for Brain Development: Andrew Jaffe, Apua Paquola, Daniel Weinberger, Jennifer Erwin, Jooheon Shin, Michael McConnell, Richard Straub, Rujuta Narurkar

Mayo Clinic: Alexej Abyzov, Taejeong Bae, Yeongjun Jang, Yifan Wang NIMH: Anjene Addington, Geetha Senthil

Sage Bionetworks: Cindy Molitor, Mette Peters

Salk Institute for Biological Studies: Fred H. Gage, Meiyan Wang, Patrick Reed, Sara Linker Stanford University: Alexander Urban, Bo Zhou, Reenal Pattni, Xiaowei Zhu

Universitat Pompeu Fabra: Aitor Serres Amero, David Juan, Inna Povolotskaya, Irene Lobon, Manuel Solis Moruno, Raquel Garcia Perez, Tomas Marques-Bonet

University of Barcelona: Eduardo Soriano University of California, Los Angeles: Gary Mathern

University of California, San Diego: Danny Antaki, Dan Averbuj, Eric Courchesne, Joseph G. Gleeson, Laurel L. Ball, Martin W. Breuss, Subhojit Roy, Xiaoxu Yang, Changuk Chung

University of Michigan: Chen Sun, Diane A. Flasch, Trenton J. Frisbie Trenton, Huira C. Kopera, Jeffrey M. Kidd, John B. Moldovan, John V. Moran, Kenneth Y. Kwan, Ryan E. Mills, Sarah B. Emery, Weichen Zhou, Xuefang Zhao

University of Virginia: Aakrosh Ratan

Yale University: Adriana Cherskov, Alexandre Jourdon, Flora M. Vaccarino, Liana Fasching, Nenad Sestan, Sirisha Pochareddy, Soraya Scuder

## References

1. Leventer, R.J., Guerrini, R. & Dobyns, W.B. Malformations of cortical development and epilepsy. Dialogues Clin Neurosci 10, 47–62 (2008).

2. Barkovich, A.J., Dobyns, W.B. & Guerrini, R. Malformations of cortical development and epilepsy. Cold Spring Harb Perspect Med 5, a022392 (2015).

3. Blumcke, I. et al. The clinicopathologic spectrum of focal cortical dysplasias: a consensus classification proposed by an ad hoc Task Force of the ILAE Diagnostic Methods Commission. Epilepsia 52, 158–74 (2011).

4. Choi, S.A. & Kim, K.J. The Surgical and Cognitive Outcomes of Focal Cortical Dysplasia. J Korean Neurosurg Soc 62, 321–327 (2019).

5. Poduri, A. et al. Somatic activation of AKT3 causes hemispheric developmental brain malformations. Neuron 74, 41–8 (2012).

6. Lee, J.H. et al. De novo somatic mutations in components of the PI3K-AKT3-mTOR pathway cause hemimegalencephaly. Nat Genet 44, 941–5 (2012).

7. Baldassari, S. et al. Dissecting the genetic basis of focal cortical dysplasia: a large cohort study. Acta Neuropathol 138, 885–900 (2019).

8. Sim, N.S. et al. Precise detection of low-level somatic mutation in resected epilepsy brain tissue. Acta Neuropathol 138, 901–912 (2019).

9. Dou, Y. et al. Accurate detection of mosaic variants in sequencing data without matched controls. Nat Biotechnol 38, 314–319 (2020).

10. Yang, X. et al. DeepMosaic: Control-independent mosaic single nucleotide variant detection using deep convolutional neural networks. bioRxiv (2021).

11. Huang, A.Y. et al. MosaicHunter: accurate detection of postzygotic single-nucleotide mosaicism through next-generation sequencing of unpaired, trio, and paired samples. Nucleic Acids Res 45, e76 (2017).

12. Wang, Y. et al. Comprehensive identification of somatic nucleotide variants in human brain tissue. Genome Biol 22, 92 (2021).

13. Breuss, M.W. et al. Autism risk in offspring can be assessed through quantification of male sperm mosaicism. Nat Med 26, 143–150 (2020).

14. Yang, X. et al. Developmental and temporal characteristics of clonal sperm mosaicism. Cell 184, 4772–4783 e15 (2021).

15. Garcia, C.A.B. et al. mTOR pathway somatic variants and the molecular pathogenesis of hemimegalencephaly. Epilepsia Open 5, 97–106 (2020).

16. Pelorosso, C. et al. Somatic double-hit in MTOR and RPS6 in hemimegalencephaly with intractable epilepsy. Hum Mol Genet 28, 3755–3765 (2019).

17. Breuss, M.W. et al. Somatic mosaicism in the mature brain reveals clonal cellular distributions during cortical development. Nature (2022).

18. Bae, T. et al. Somatic mutations reveal hypermutable brains and are associated with neuropsychiatric disorders. medRxiv (2022).

19. Bozic, I. et al. Accumulation of driver and passenger mutations during tumor progression. Proc Natl Acad Sci U S A 107, 18545–50 (2010).

20. Alexandrov, L.B. et al. The repertoire of mutational signatures in human cancer. Nature 578, 94–101 (2020).

21. Kim, M. & Costello, J. DNA methylation: an epigenetic mark of cellular memory. Exp Mol Med 49, e322 (2017).

22. Szklarczyk, D. et al. STRING v11: protein-protein association networks with increased coverage, supporting functional discovery in genome-wide experimental datasets. Nucleic Acids Res 47, D607–D613 (2019).

23. Bedrosian, T.A. et al. Detection of brain somatic variation in epilepsy-associated developmental lesions. medRxiv (2021).

24. Lai, D. et al. Somatic mutation involving diverse genes leads to a spectrum of focal cortical malformations. medRxiv (2021).

25. Tarabeux, J. et al. Rare mutations in N-methyl-D-aspartate glutamate receptors in autism spectrum disorders and schizophrenia. Transl Psychiatry 1, e55 (2011).

26. Bezprozvanny, I. Calcium signaling and neurodegenerative diseases. Trends Mol Med 15, 89–100 (2009).

27. Su, M.Y. et al. Hybrid Structure of the RagA/C-Ragulator mTORC1 Activation Complex. Mol Cell 68, 835–846 e3 (2017).

28. Chen, J. et al. KLHL22 activates amino-acid-dependent mTORC1 signalling to promote tumorigenesis and ageing. Nature 557, 585–589 (2018).

29. Behar, T.N. et al. Glutamate acting at NMDA receptors stimulates embryonic cortical neuronal migration. J Neurosci 19, 4449–61 (1999).

30. Paoletti, P., Bellone, C. & Zhou, Q. NMDA receptor subunit diversity: impact on receptor properties, synaptic plasticity and disease. Nat Rev Neurosci 14, 383–400 (2013).

31. Strehlow, V. et al. GRIN2A-related disorders: genotype and functional consequence predict phenotype. Brain 142, 80–92 (2019).

32. Prickett, T.D. & Samuels, Y. Molecular pathways: dysregulated glutamatergic signaling pathways in cancer. Clin Cancer Res 18, 4240–6 (2012).

33. Ruvinsky, I. & Meyuhas, O. Ribosomal protein S6 phosphorylation: from protein synthesis to cell size. Trends Biochem Sci 31, 342–8 (2006).

34. Sumimoto, H., Imabayashi, F., Iwata, T. & Kawakami, Y. The BRAF-MAPK signaling pathway is essential for cancer-immune evasion in human melanoma cells. J Exp Med 203, 1651–6 (2006).

35. Ornitz, D.M. & Itoh, N. The Fibroblast Growth Factor signaling pathway. Wiley Interdiscip Rev Dev Biol 4, 215–66 (2015).

36. Chen, X. et al. TNF-alpha-Induced NOD2 and RIP2 Contribute to the Up-Regulation of Cytokines Induced by MDP in Monocytic THP-1 Cells. J Cell Biochem 119, 5072–5081 (2018).

37. Yates, T.M. et al. SLC35A2-related congenital disorder of glycosylation: Defining the phenotype. Eur J Paediatr Neurol 22, 1095–1102 (2018).

38. Paganini, C. et al. Calcium activated nucleotidase 1 (CANT1) is critical for glycosaminoglycan biosynthesis in cartilage and endochondral ossification. Matrix Biol 81, 70–90 (2019).

39. Lee, N. et al. Neuronal migration disorders: positron emission tomography correlations. Ann Neurol 35, 290–7 (1994).

40. Kim, Y.H. et al. Neuroimaging in identifying focal cortical dysplasia and prognostic factors in pediatric and adolescent epilepsy surgery. Epilepsia 52, 722–7 (2011).

41. Nowakowski, T.J. et al. Spatiotemporal gene expression trajectories reveal developmental hierarchies of the human cortex. Science 358, 1318–1323 (2017).

42. Coe, B.P. et al. Neurodevelopmental disease genes implicated by de novo mutation and copy number variation morbidity. Nat Genet 51, 106–116 (2019).

43. Ridley, A.J. et al. Cell migration: integrating signals from front to back. Science 302, 1704–9 (2003).

44. Brini, M., Cali, T., Ottolini, D. & Carafoli, E. Neuronal calcium signaling: function and dysfunction. Cell Mol Life Sci 71, 2787–814 (2014).

45. Lamparello, P. et al. Developmental lineage of cell types in cortical dysplasia with balloon cells. Brain 130, 2267–76 (2007).

46. Englund, C., Folkerth, R.D., Born, D., Lacy, J.M. & Hevner, R.F. Aberrant neuronal-glial differentiation in Taylor-type focal cortical dysplasia (type IIA/B). Acta Neuropathol 109, 519–33 (2005).

47. Kim, S. et al. Strelka2: fast and accurate calling of germline and somatic variants. Nat Methods 15, 591–594 (2018).

48. Benjamin, D. et al. Calling Somatic SNVs and Indels with Mutect2. bioRxiv (2019).

49. Yang, X. et al. Genomic mosaicism in paternal sperm and multiple parental tissues in a Dravet syndrome cohort. Sci Rep 7, 15677 (2017).

50. Untergasser, A. et al. Primer3Plus, an enhanced web interface to Primer3. Nucleic Acids Res 35, W71–4 (2007).

51. Untergasser, A. et al. Primer3--new capabilities and interfaces. Nucleic Acids Res 40, e115 (2012).

52. Lee, J. et al. Mutalisk: a web-based somatic MUTation AnaLyIS toolKit for genomic, transcriptional and epigenomic signatures. Nucleic Acids Res 46, W102–W108 (2018).

53. Bindea, G. et al. ClueGO: a Cytoscape plug-in to decipher functionally grouped gene ontology and pathway annotation networks. Bioinformatics 25, 1091–3 (2009).

54. Koizumi, H., Tanaka, T. & Gleeson, J.G. Doublecortin-like kinase functions with doublecortin to mediate fiber tract decussation and neuronal migration. Neuron 49, 55–66 (2006).

55. Wang, F. et al. RNAscope: a novel in situ RNA analysis platform for formalin-fixed, paraffin-embedded tissues. J Mol Diagn 14, 22–9 (2012).

